# Collective amoeboid dynamics drives colonization of drug-resistant ovarian cancer cells

**DOI:** 10.1101/2024.10.05.616763

**Authors:** Shyamili Goutham, Anchita Gopikrishnan, BV Harshavardhan, Nivedhya Venas, Radhakrishnan Sabarinathan, Mohit Kumar Jolly, Ramray Bhat

**Affiliations:** Department of Developmental Biology and Genetics, Indian Institute of Science, Bengaluru, India, 560012; Department of Bioengineering, Indian Institute of Science, Bengaluru, India, 560012; National Centre for Biological Sciences, Tata Institute of Fundamental Research, Bengaluru, India

## Abstract

Epithelial ovarian cancer (EOC) is characterized by resistance to platinum-based therapy, resulting in rapid progression and poor survival. Here, we ask whether drug resistance and invasiveness coevolve to drive metastasis. Selection experiments involving pulsed carboplatin exposure established isogenic chemoresistant variants of lines, which typify high-grade serous ovarian carcinoma (HGSOC), the most aggressive type of EOC. Time-lapse imaging showed enhanced migration of resistant single cells and their collectives. Resistant cell spheroids spread faster on Collagen I substrata than sensitive controls. The resistant OVCAR-3 transcriptome was ontologically enriched for migration and showed overlap with previously reported markers of resistance in EOC patients and other evolved lines. Gene set enrichment predicted transition between epithelial, mesenchymal, and amoeboid states is higher in resistance compared to control lines. Lower matrix adhesion, weak focal adhesion, and highly deformable and translatory dynamics of cell collectives indicated that resistant cancer cells displayed a unique collective amoeboid-like migration. When injected intraperitoneally into immunodeficient mice, resistant cells colonized to a greater extent on parietal mucosae. Ex vivo, suspended resistant cells formed moruloids associated with quicker peritoneal adhesion, clearing human coelomic mesothelial monolayers with higher efficiency. Knockdown in resistant OVCAR-3 cells of two upregulated proteins, E-cadherin and LGALS3BP, had distinct consequences. E-cadherin knockdown partially restored sensitivity to carboplatin but did not affect invasion. In contrast, silencing LGALS3BP decreased invasion but not resistance. Our results suggest that drug resistance and invasiveness could coevolve through the upregulation of distinct trait drivers in EOC.

## Introduction

Despite impressive strides made in understanding the mechanisms behind the progression of epithelial ovarian cancer (EOC), the mortality rate associated with the disease remains alarmingly high (Coleman et al., 2011; Siegel et al., 2021). Upon diagnosis, ovarian cancer is managed through a combination of surgical cytoreduction and treatments with chemotherapeutics (Ortiz et al., 2022) and, in some cases, personalized medicine (Morand et al., 2021). Patients afflicted with EOC, especially its aggressive variant known as high-grade serous ovarian cancer (HGSOC), initially respond well to a combination of platinum and taxane drugs (Lheureux et al., 2019). However, most who do show a good response eventually relapse and develop a chemoresistant form of the disease, which contributes to high patient mortality (Guo et al., 2022).

Several studies have sought to address how resistance evolves in ovarian tumor populations. Platinum-based chemo drugs induce apoptosis by adducing themselves to purine bases of DNA and triggering their damage (Eastman, 1987; Rabik & Dolan, 2007). Therefore, cells become resistant by efficiently mediating drug efflux or improving their ability to repair nucleic acid damage (Cornelison et al., 2017). Repeated rounds of drug exposure in tumor cell populations increase the frequency of resistant variants that possess unique genomic alterations and distinct transcriptomic and proteomic profiles relative to their sensitive counterparts. A recent elegant study on paraffin-embedded tissue sections from HGSOC patients receiving neoadjuvant chemotherapy revealed transcriptional upregulation of the AP-1 transcription factor family genes and copy number amplification of a chromosomal segment that subsumes the SIK2; in combination with platinum-based therapy, inhibition of both synergistically decrease resistant cell viability (Javellana et al., 2022). Such transcriptome-wide alterations also result in modifications in other phenotypic traits that sculpt cancer progression. For example, overexpression of miR-296-3p has been shown to significantly enhance the proliferation and migration of ovarian cancer cells in vitro, showing a greater degree of drug resistance (Sun et al., 2024). Transcriptomic changes associated with drug resistance may also result in altered matrisomal dynamics that can, in turn, modify the interaction of cancer cells with their tissue microenvironment (Salemme et al., 2023). A recent study uncovers the upregulation in resistant cells of the extracellular matrix protein Periostin, which contributes to recurrence in EOC cells by enhancing stemness (Huang et al., 2023). Acquisition of drug resistance is also often correlated with morphological-molecular changes that signify a transition from the epithelial state into a more mesenchymal one (EMT) (Kajiyama et al., 2007; Rohnalter et al., 2015). The link between EMT and the evolution of drug resistance has been proposed to be established by nodal proteins such as those belonging to the Snail family (Kielbik et al., 2023). Mediators of EMT represent novel druggable targets in chemo-refractory cancers (Tangsiri et al., 2024).

Apart from the epithelial and mesenchymal states, recent studies suggest the transitioning of mesenchymal cells to an amoeboid state as an adaptive measure to bypass microenvironmental stress, which may also include the induction of drug resistance mechanisms (Emad et al., 2020; Ketchen et al., 2021). Amoeboid migration, typically characterized for immune cells and dictyostelid amoebae, achieves motility without relying on strong substrate adhesion, as is seen for mesenchymal cells (Barry & Bretscher, 2010; George et al., 2023). As a result, amoeboid migration, which is characterized by short-lived actomyosin-rich pseudopodia, is faster than mesenchymal migration (Lämmermann et al., 2008) and has been proposed to contribute to metastatic kinetics (Crosas-Molist et al., 2017; Driscoll et al., 2024). These studies notwithstanding, formal analyses of whether and how phenotypic traits tend to co-evolve with drug resistance, and if such coevolution has consequences on HGSOC progression in a drug-agnostic context are few and far between.

In the current study, we develop an experimental model of platinum chemoresistance by exposing the HGSOC lines OVCAR-3 and COV362 to periodic pulses of the drug carboplatin. Having ascertained the evolution of resistance, we rigorously probe its migratory potential and show how the resistance is associated not just with enhanced single-cell motility but also collective migration. RNA sequencing followed by ontological analysis surprisingly suggests the evolution of a complex hybrid state that possesses signatures of epithelial, mesenchymal, and amoeboid state, which is consistent with their collective cellular behavior, low ECM adhesion, upregulation of metastasis-related extracellular matrix genes, and in vivo peritoneal colonization morphologies.

## Results

### Carboplatin-resistant cells migrate faster than sensitive controls

Carboplatin-resistant variants of the HGSOC lines OVCAR-3 and COV362 were developed by repeated exposure to 10-15 cycles of IC_50_ dose of carboplatin (refer to Figures 1A and the methods subsection titled “establishing resistant cell lines”), after which their metabolic viability was assessed using resazurin assay (Figure 1B). Carboplatin IC_50_ for the parental OVCAR-3 line was found to be 21.77 μM ± 0.96, whereas for the resistant OVCAR-3 or resOVCAR-3, the IC_50_ was found to be 70.90 μM ± 3.77 (Figure 1C; p <0.0001 significance established using unpaired student’s t-test). IC_50_ values for the parental COV362 and resistant COV362 lines were also estimated to be 50.18 μM ± 13.42 and 118.1 μM ± 12.77, respectively (Figure S1A, Figure S1B; p = 0.01 significance established using unpaired student’s t-test). Similarly, IC_50_ for a second independently evolved resistant OVCAR-3 (resOVCAR-3(2)) line was estimated to be 62.21 μM ± 1.66 (Figure S2A, Figure S2B; p = 0.02 significance established using unpaired student’s t-test). To assess whether the differences in values within resazurin-based assays were due to effects on cell viability or metabolic activity, OVCAR-3 and resOVCAR-3 were treated for 72 h with 30 μM carboplatin and their numbers calculated using hemocytometry: resOVCAR3 cells showed a 4.3-fold higher cell number indicating greater cell viability in the presence of carboplatin (Figure 1D; p < 0.01 significance established using unpaired student’s t-test).

**Figure 1:**
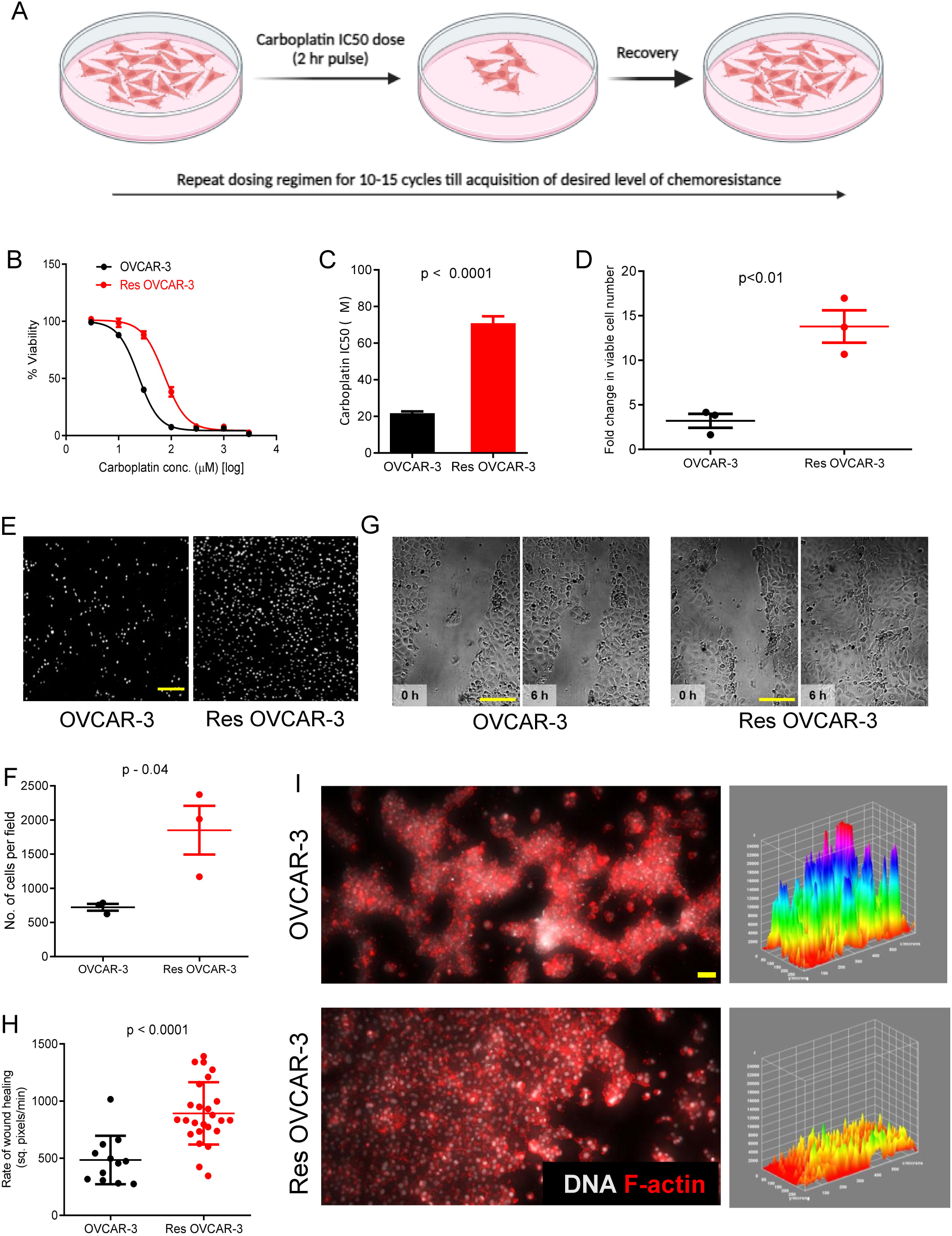
Carboplatin-resistant cells show enhanced migratory properties. (A) Schematic showing the pulsing regimen used for the generation of carboplatin-resistant cell lines. Illustration made using BioRender. (B, C) Graphs depicting viability of OVCAR-3 and resOVCAR-3 at different concentrations of carboplatin evaluated using resazurin assay. Each point in the viability curves and bars indicates mean ± SEM. n≥5 for all viability assays performed. Significance was measured by an unpaired student’s t-test. (D) Graph showing the viability of OVCAR-3 and resOVCAR-3 upon 30 µM carboplatin treatment evaluated by estimating the viable number of cells using a hemocytometer. (E, F) Invasion of OVCAR-3 and resOVCAR-3 cells through Collagen-I coated cell culture inserts across a serum concentration gradient. (E) Epifluorescent micrographs of OVCAR-3 (left) and resOVCAR-3 (right) cells stained with propidium iodide (PI) for DNA (white) on the lower side of the transwell insert. Scale bar - 500 µm. (F) Graph showing the number of cells per field that have invaded through Collagen-I coated cell culture inserts. (G, H) Scratch assay for OVCAR-3 and resOVCAR-3 cells measured through rate of scratch disappearance (refer to videos S1, S2). (G) Phase contrast images of scratches on OVCAR-3 and resOVCAR-3 cellular monolayers at 0 h and 6 h. Scale bar - 200 µm. (H) Graph showing the rate of disappearance of scratch through collective cell migration. (I) Colonization (adhesion and spread) of OVCAR-3 and resOVCAR-3 spheroids on top of 1 mg/mL Collagen-I scaffolds in 72 h (refer to videos S3, S4). Fluorescent confocal micrographs of OVCAR-3 (top left) and resOVCAR-3 (bottom left) clusters on top of 1 mg/mL Collagen-I stained for F-actin (red using phalloidin) and DNA (white using DAPI). Scale bar – 50 µm. Corresponding images on the right show 3D representations of the attached clusters heat-colored to represent their height. Experiments performed n ≥ 3 times. Bars represent mean ± SEM. Significance is measured using an unpaired t-test.

Drug-resistant cancers are usually aggressive in their progression with increased incidence of metastases (Norouzi et al., 2018). Accordingly, we sought to test whether resOVCAR-3 cells invade faster through an extracellular matrix (ECM) milieu with respect to their control counterparts. Transwell migration assays through Collagen-I gels showed greater single-cell invasion for resOVCAR-3 cells compared with controls (Figure 1E, epifluorescent micrographs of OVCAR-3 (left) and resOVCAR-3 (right) cells stained with propidium iodide (PI) for DNA (white against black background)); Figure 1F, a 2.6-fold increase in mean cell number per field for resOVCAR-3 cells, p=0.04 with significance calculated using an unpaired student’s t-test), resCOV362 cells (Figure S3A, epifluorescent micrographs of COV362 (left) and resCOV362 (right) cells stained with PI for DNA (white against black background)); Figure S3B, 4.4-fold increase in mean cell number per field for resCOV362 cells; p=0.05 with significance calculated using an unpaired student’s t-test) and the second independent resOVCAR-3(2) line (Figure S4A, epifluorescent micrographs of OVCAR-3 (left) and resOVCAR-3(2) (right) cells stained with PI for DNA (white against black background)); Figure S4B, 6.9-fold increase in mean cell number per field for resOVCAR-3(2) cells; p=0.04 with significance calculated using an unpaired student’s t-test).

In order to examine the collective migration, scratches were made to confluent cell monolayers, and the rate of scratch disappearance due to cell movement was examined. We observed that the resOVCAR-3 cell sheets moved faster than control OVCAR-3 cells (Figure 1G: phase contrast micrographs of OVCAR-3 (left top 0h and left bottom 6h) and resOVCAR-3 (right top 0h and right bottom 6h), Figure 1H: 1.8-fold increase in the rate of scratch disappearance for resOVCAR-3 cells p<0.0001 with significance calculated using an unpaired student’s t-test, videos S1, S2: time-lapse videos showing scratch disappearance in OVCAR-3 and resOVCAR-3 cell layers). In addition, suspended spheroids of resOVCAR-3 cells attached and spread faster on 1 mg/ml Collagen I scaffolds compared with OVCAR-3 (Figure 1I, fluorescent confocal micrographs of sensitive control (top) and resOVCAR-3 cells (bottom) with signals for F-actin (red using phalloidin) and DNA (white using DAPI. Refer to videos S3, S4). Corresponding images on the right show 3D representations of the attached clusters heat-colored to represent their height, showing greater dispersion and flattening of resOVCAR-3 aggregates resulting in smaller peak heights.

### resOVCAR-3 shows transcriptomic signatures cognate with migration

We next sought to probe the difference in gene expression due to the acquisition of resistance to carboplatin. RNA sequencing was performed with three independent passages of control and resOVCAR-3 cells (Figure S5 shows the principal component analysis showing the proximity of overall cell states of the replicates). Differential gene expression analysis between the resistant and sensitive cell lines showed that 1457 genes were upregulated, while 1376 genes were downregulated in resistant compared to sensitive cells, with |log-fold change| > 1 and FDR (q-value) < 0.05 (Figure 2A). Gene ontological analysis for the significantly up- and down-regulated expressions of genes for biological functions predicted mRNA alterations associated with cell motility and morphogenesis (Figure 2B top). This was further confirmed through gene set enrichment based on the KEGG pathways, which, in addition to altered motility and cell-ECM interaction, implicated an altered regulation of the PI3K-AKT pathway (Figure 2B bottom). Transduction through this signaling pathway (through a cognate signature set) has been shown to be associated with poor survival, harsher progression, and poorer response to therapy in patients with serous ovarian cancer (Carvalho et al., 2022). We used patient-derived gene signatures for poor prognosis, which comprised 20 genes, to assay if the transcriptome of resOVCAR-3 recapitulates the transcriptome of the patients: of the 20 genes (related to the PI3K pathways), 12 (COL1A1, LAMA5, LAMC1, ITGB1, COL4A2, EFNA1, EPHA2, LAMC2, ITGA6, ITGA3, ITGB4, FN1) showed an upregulation in our resistant cell line, and one gene, IGF1, was not available in our RNA sequencing data (Figure 2C; refer to Table S1 for log2fold change and p values).

**Figure 2:**
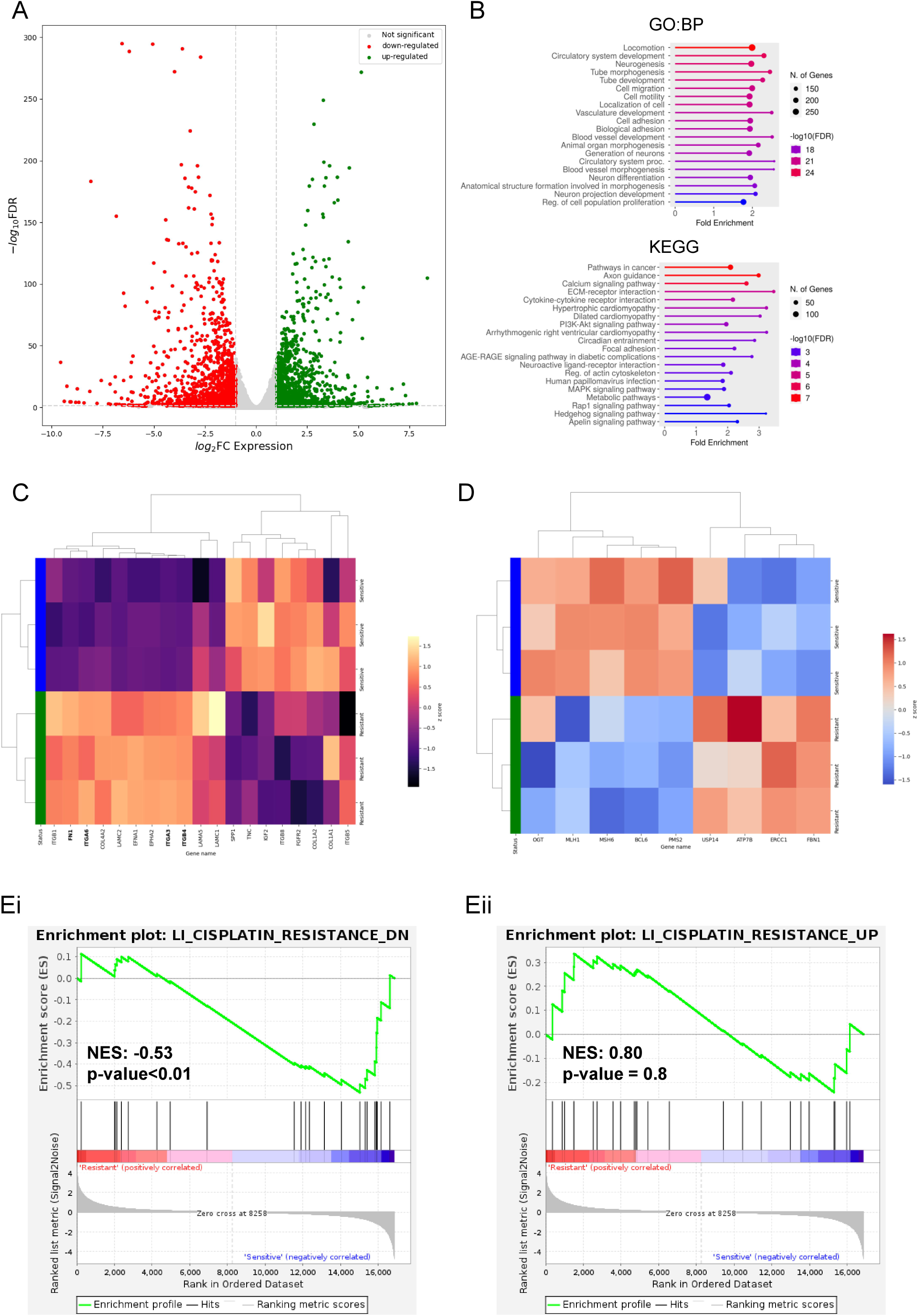
Transcriptomic profiling reveals signatures associated with migration in resOVCAR-3. (A) Volcano plot depicts the differentially expressed genes in resOVCAR-3 with respect to OVCAR-3. Each dot represents a gene. The x-axis represents the log2 fold change in expression values in resOVCAR-3 versus OVCAR-3, and the y-axis shows the −log 10 adjusted p-value. The vertical and horizontal dashed lines indicate the threshold of FDR (q-value) <0.05 and │log2fold change (expression)│ of 1. (B) Gene ontology (biological processes) analysis plots (FDR cutoff – 0.05) (top) and KEGG enrichment analysis plots of OVCAR-3 and resOVCAR-3 transcriptomes (FDR cutoff – 0.05) (bottom) generated using ShinyGO (Ge et al., 2020). (C) Transcriptional changes of 19 genes (Carvalho et al., 2022) associated with poor survival in serous ovarian cancer are represented as z-scores with respect to OVCAR-3 (green rows) and res OVCAR-3 (blue rows). The genes marked in bold are significantly upregulated in resOVCAR-3. (D) Transcriptional changes of 10 genes, chosen based on their relevance in chemotherapy resistance mechanisms (Du et al., 2016; Kalayda et al., 2012; Mangala et al., 2009; Ortiz et al., 2022; J. Shen et al., 2020; L. Shen et al., 2021; Wang et al., 2022; Zhao et al., 2018; Zhou et al., 2018), represented as z-scores with respect to OVCAR-3 (green rows) and resOVCAR-3 (blue rows). (Ei, Eii) Gene set enrichment analysis of resOVCAR-3 compared to OVCAR-3. Gene sets related to cisplatin resistance in ovarian cancer cell lines were used (Li et al., 2007; Liberzon et al., 2011).

We also examined the expression of a second curated set of 10 genes chosen based on their dysregulation in drug-resistant ovarian cancer progression (Ortiz et al., 2022). This set included the genes involved in influx (CTR1) and efflux (ATP7B) (Kalayda et al., 2012; Mangala et al., 2009) of carboplatin. CTR1 did not appear in our sequencing analysis. The efflux transporter gene ATP7B was upregulated in resOVCAR-3 cells, consistent with previous literature on its expression in drug-resistant cells (Figure 2D). The gene set also included DNA mismatch repair genes, which are generally mutated/silenced in EOC lines (Milanesio et al., 2020; Ovejero-Sánchez et al., 2023). As expected, we found a corresponding downregulation of MSH6, MLH1, and PMS2, in resOVCAR-3 cell line replicates. Additionally, Zhao and co-workers have shown that high levels of expression for these genes are associated with favorable prognosis post-platinum therapy (Zhao et al., 2018). resOVCAR3 cells also showed an upregulation for the nucleotide excision repair protein coding gene ERCC1, which is known to be elevated in cisplatin-resistant cells (Du et al., 2016). We further added another pair of genes - USP14 and BCL6- to the gene set based on previous reports, which showed their elevation with platinum resistance (J. Shen et al., 2020). Our expression analysis of resOVCAR3 cells showed an upregulation in the case of USP14 but a downregulation in BCL6. Similarly, the misexpression of two other genes in the gene set, the enzyme O-GlcNAc transferase (OGT) (Zhou et al., 2018) and the extracellular matrix protein Fibrillin-1 (FBN1) (Wang et al., 2022), is recapitulated by the resOVCAR-3 (refer to Table S2 for log2fold change and p values). However, the expression for PGC1/PPARGC1A was lower in resOVCAR-3 cells, unlike what has been observed in the cisplatin resistance (L. Shen et al., 2021); this could be due to minor differences in metabolic responses of cells to carboplatin and cisplatin.

Furthermore, we obtained expression signatures from cisplatin-resistant ovarian cancer line (ACRP) from MsigDB (Li et al., 2007; Liberzon et al., 2011), which showed similar trends with the resOVCAR-3 transcriptome, particularly for the downregulated set of genes (Figure 2Ei and ii).

### Drug-resistant cells show stronger signatures of migration

Our observations of the migratory behaviors of resOVCAR-3 cells combined with results from gene ontology analyses motivated us to examine the possibility of the transition of resistant cells to a more mesenchymal expression state. This is in consonance with a strong association in ovarian and other cancers of resistance evolution with an epithelial-to-mesenchymal transition (EMT) (Debaugnies et al., 2023; Kralj et al., 2023). We performed single-sample Gene Set Enrichment Analysis (ssGSEA) on the control OVCAR-3 and resOVCAR-3 transcriptomes and observed higher scores for both epithelial and mesenchymal marker signatures in resOVCAR-3 cells, which suggested that drug resistance may be associated with the acquisition of a hybrid phenotype (Figure 3A, B; p = 0.002 and 0.06; significance established using unpaired student’s t-test). A hybrid or partial EMT state is intermediate within the spectrum characterized by the expression of both epithelial and mesenchymal markers, resulting in phenotypic characteristics more aggressive than the complete transited mesenchymal state (Brown et al., 2022; Kisoda et al., 2022). The existence of resOVCAR-3 cells in a hybrid EMT state was further indicated by the high partial EMT score obtained using ssGSEA analysis (Figure 3C, p = 0.002; significance established by unpaired student’s t-test). Results from the bioinformatics analyses were further confirmed using real-time quantitative PCR, wherein mRNA levels of the epithelial marker E-cadherin (CDH1) and the mesenchymal marker fibronectin-1 (FN1) were found to be 2.5-fold and 2.8-fold higher in resOVCAR-3 cells compared with control OVCAR-3 (Figure 3D, E; p<0.0001 and p=0.01 respectively; significance established using unpaired student’s t-test). CDH1 and FN1 protein levels were also found to be upregulated immunocytochemically in resOVCAR-3 compared to OVCAR-3 cells (Figure 3F, G; green: secondary antibody, red: F-actin using phalloidin and white: DNA using DAPI; see Figure S6, S7 for no primary antibody control).

**Figure 3:**
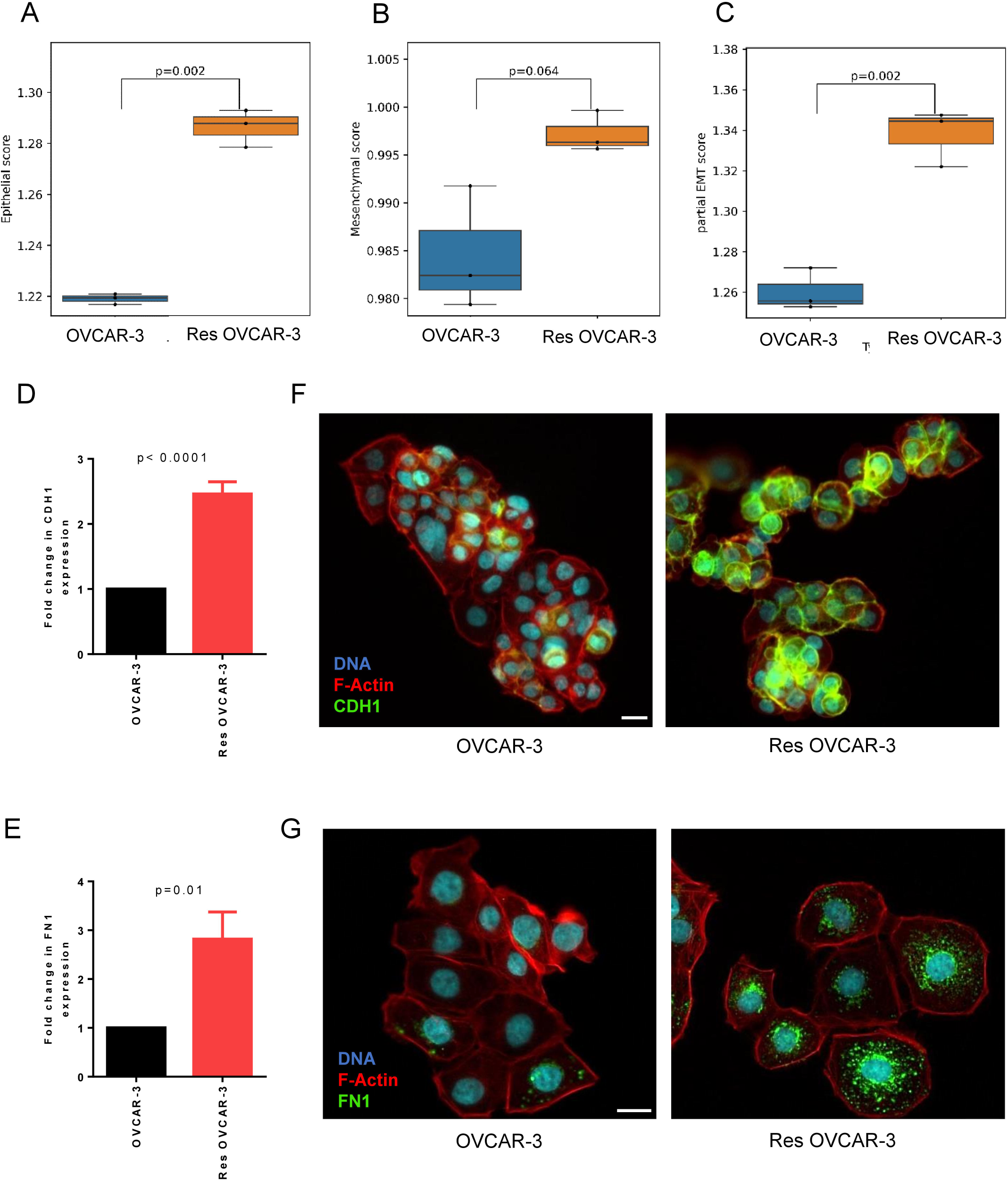
ResOVCAR-3 cells show enrichment for both epithelial and mesenchymal state signatures. (A, B, C) ssGSEA analysis plots from the RNA sequencing data presented in Figure 2 indicate the expression profile of curated epithelial (A) and mesenchymal (B) markers in OVCAR-3 and resOVCAR-3 cells. (C) ssGSEA plots depicting the expression of partial EMT signatures in OVCAR-3 and resOVCAR-3. (D, E) Graph showing elevated mRNA levels of CDH1 (D) and FN1 (E) in resOVCAR-3 compared to OVCAR-3 (18sRNA used as internal control) observed using qPCR analysis. (F, G) Fluorescent confocal micrographs of OVCAR-3 and resOVCAR-3 cells cultured as monolayers stained for E-cadherin (F) and fibronectin (G) (green), counterstained with F-actin (phalloidin; red) and DNA (DAPI; white) (refer to Figures S6 and S7 for no primary antibody control). Experiments performed n ≥ 3 times. Scale bar – 20 μm.

Motility in partially and fully transited mesenchymal cells is mediated through high adhesion of cellular protrusion to underlying extracellular matrix substrata. Therefore, adhesion to Collagen I substrata was compared between suspended OVCAR-3 and resOVCAR-3 cells: in the same time interval, a lesser number of the latter adhered compared to sensitive control cells (Figure 4A, epifluorescent micrographs of OVCAR-3 (left) and resOVCAR-3 (right) cells stained with PI for DNA (white against black background); Figure 4B, 1.9-fold difference in mean cell number per field; p = 0.001 with significance calculated using an unpaired student’s t-test). We hypothesized that the resistant cells may employ an amoeboid state, wherein cells achieve higher velocities by eschewing strong cell-ECM adhesion (Graziani et al., 2022). Cancer cells are known to transit from mesenchymal to amoeboid modes (MAT) partially or completely (B. Huang et al., 2014; Paňková et al., 2010; Talkenberger et al., 2017), and a new signature inclusive of EMT and MAT has been proposed (EMAT) (Emad et al., 2020) to also account for such hybrid states. To understand if there is a significant shift in the population of cells exhibiting amoeboid signatures, ssGSEA analysis was performed on transcriptomes of OVCAR-3 and resOVCAR-3 cells. Although statistically insignificant, our results indicate increased MAT signatures in resOVCAR-3 cell populations (Figure S8). However, further ssGSEA analysis to study the expression profile of genes involved in the epithelial-mesenchymal-amoeboid transitions (EMAT) yielded a significant upregulation of these markers in the res-OVCAR3 cell populations (Figure 4C; p = 0.01 with significance calculated using an unpaired student’s t-test).

**Figure 4:**
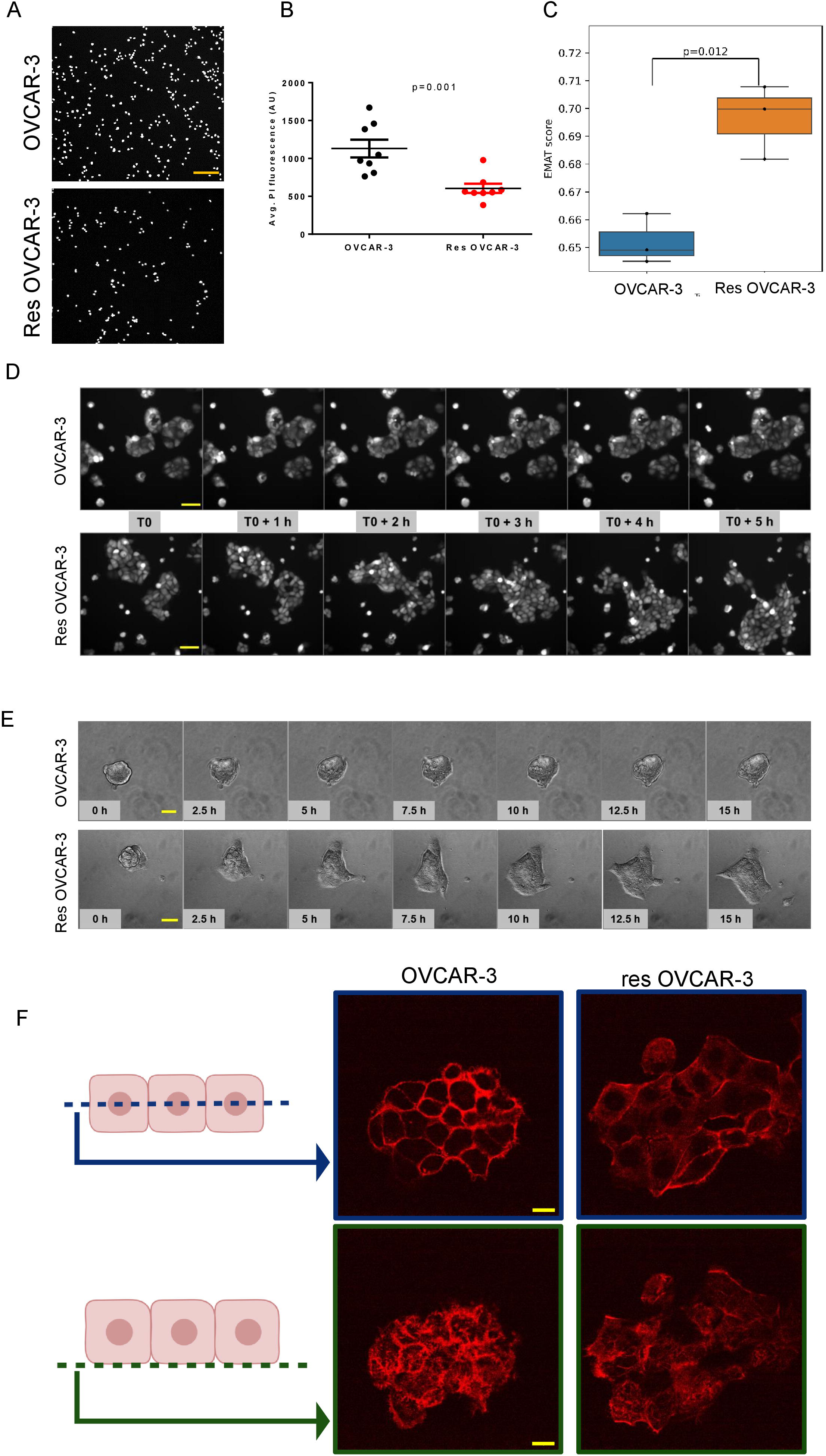
ResOVCAR-3 cells exhibit a collective amoeboid phenotype. (A, B) Adhesion assay was performed by seeding cells on Collagen I, followed by fixing and staining adhered cells with propidium iodide (PI) to quantify differences in adhesion potential. (A) Representative epifluorescence micrographs showing number of adhered cells per field in resOVCAR-3 samples compared to OVCAR-3. Scale bar – 200 μm. (B) The graph shows net PI fluorescence indicative of the number of adhered cells on Collagen-I. (C) ssGSEA analysis plot depicting the expression of markers of epithelial-mesenchymal-amoeboid transition (EMAT) states in OVCAR-3 and resOVCAR-3 cell populations. (D) Epifluorescence micrographs captured using time-lapse imaging showing migratory behaviors of a patch of fluorescently labeled OVCAR-3 (top) and resOVCAR-3 (bottom) cells cultured in normal 2D conditions. Scale bar – 100 μm. Refer to videos S5 and S6 for the same. (E) Bright-field photomicrographs captured using time-lapse imaging showing adhesion and collective migratory properties of OVCAR-3 (top) and resOVCAR-3 (bottom) spheroids on Collagen-I. Scale bar – 50 μm. Refer to videos S3 and S4 for the same. (F) Confocal fluorescent micrographs of OVCAR-3 (left) and resOVCAR-3 (right) cellular patches stained for F-actin using phalloidin (red). The illustration on the left indicates the cellular planes imaged and represented on the right. Images on the top represent planes showing intracellular cortical F-actin signals, while images at the bottom represent basal actin stress fiber staining. Scale bar – 20 μm. Experiments performed n ≥ 3 times. Bars indicate mean ± SEM. Significance is measured using an unpaired student’s t-test.

To examine these dynamics further, we performed time-lapse fluorescence imaging on GFP/RFP expressing OVCAR-3 and resOVCAR-3 cells (Video S5, S6). To our surprise, small multicellular patches of resOVCAR-3 cells acted like giant amoebae by constantly altering their shape, sending out protrusions, and even translating en masse. In contrast, control OVCAR-3 patches grew centrifugally through cell division without net translation (Figure 4D, epifluorescence micrographs captured over specific time intervals showing migratory behaviors of a patch of fluorescently labeled OVCAR-3 (top) and resOVCAR-3 (bottom) cells cultured in normal 2D conditions).

Similar collective migratory properties of the resOVCAR-3 cells compared to control OVCAR-3 cells were also observed when 24-hr spheroids were allowed to adhere and grow on top of 1 mg/mL Collagen I scaffolds (Figure 4E, bright field photomicrographs captured over specific time intervals showing adhesion and collective migratory properties of OVCAR-3 (top) and resOVCAR-3 (bottom) clusters on Collagen-I, videos S3, S4: time-lapse videos of the same). Compared to the relatively sedate sensitive OVCAR-3 spheroids, resOVCAR-3 spheroids showed rapid en masse motility, throwing mesoscale multicellular projections and retractions. Staining such patches for F-actin (using phalloidin) and DNA (using DAPI), we observed faint supracellular actin staining in resOVCAR-3 patches with decreased intracellular cortical F-actin signals. In contrast, sensitive controls showed strong intracellular cortical staining for F-actin within patches and little sign of supracellular organization (Figure 4F, confocal micrographs depicting F-actin (red) and DNA (blue) staining of OVCAR-3 (left) and resOVCAR-3 (right) cellular patches). In addition, sensitive OVCAR-3 and resistant resOVCAR-3 cells showed strong and weak basal actin stress fiber staining (right images), respectively.

### Chemoresistant ovarian cancer cells colonize parietal peritoneal surfaces faster than sensitive cells

Having characterized the chemoresistance phenotype and the emergent migratory properties of the resistant cells, we proceeded to investigate these characteristics in the progression of aggressive ovarian cancers *in vivo*. Immunodeficient mice were injected with fluorescently labeled OVCAR-3 (GFP) and resOVCAR-3 (RFP) cells via intraperitoneal injections. Parietal peritoneal sections were surgically dissected from the mice 72 hours post-injection and imaged to detect fluorescence signals on the parietal serosa, indicating the presence of adhered and spread cancer cells (Figure 5A). Fluorescence micrographs of the peritoneum sections reveal larger areas of the tissue colonized by resOVCAR-3 cells (red, right) compared to OVCAR-3 cells(green, left) (Figure 5B, epifluorescent micrographs of stitched fields of dissected parietal serosa (boundary in purple) showing cognate fluorescence; Figure 5C shows 1.6-fold difference between the attachment of resOVCAR-3 cells; p = 0.03 as measured using paired student’s t-test). These findings were consistent with the greater spread of resistant cancer cells observed on Collagen I scaffolds. However, since peritoneal surfaces also consist of mesothelia, we strove to investigate potential differences in the interactions between suspended spheroids of control and resOVCAR-3 cells and adhered layers of untransformed human coelomic mesothelial cells (Met-5A) using time-lapse microscopic imaging (videos S7, S8). It was observed that the resOVCAR-3 cell clusters settled and spread at a faster rate amidst the mesothelial cells by clearing them, compared to OVCAR-3 cell clusters (Figure 5D epifluorescence micrographs captured over specific time intervals showing adhesion and spread of OVCAR-3 GFP (left) and resOVCAR-3 RFP (right) spheroids on Met-5A cell layer; Figure 5E shows 3.9-fold difference in spread of resOVCAR-3 cells compared with control, p = 0.002 measured using an unpaired student’s t-test). This indicates that resistant ovarian cancer cells can colonize cellular and acellular matrix surfaces more efficiently than their sensitive counterparts.

**Figure 5:**
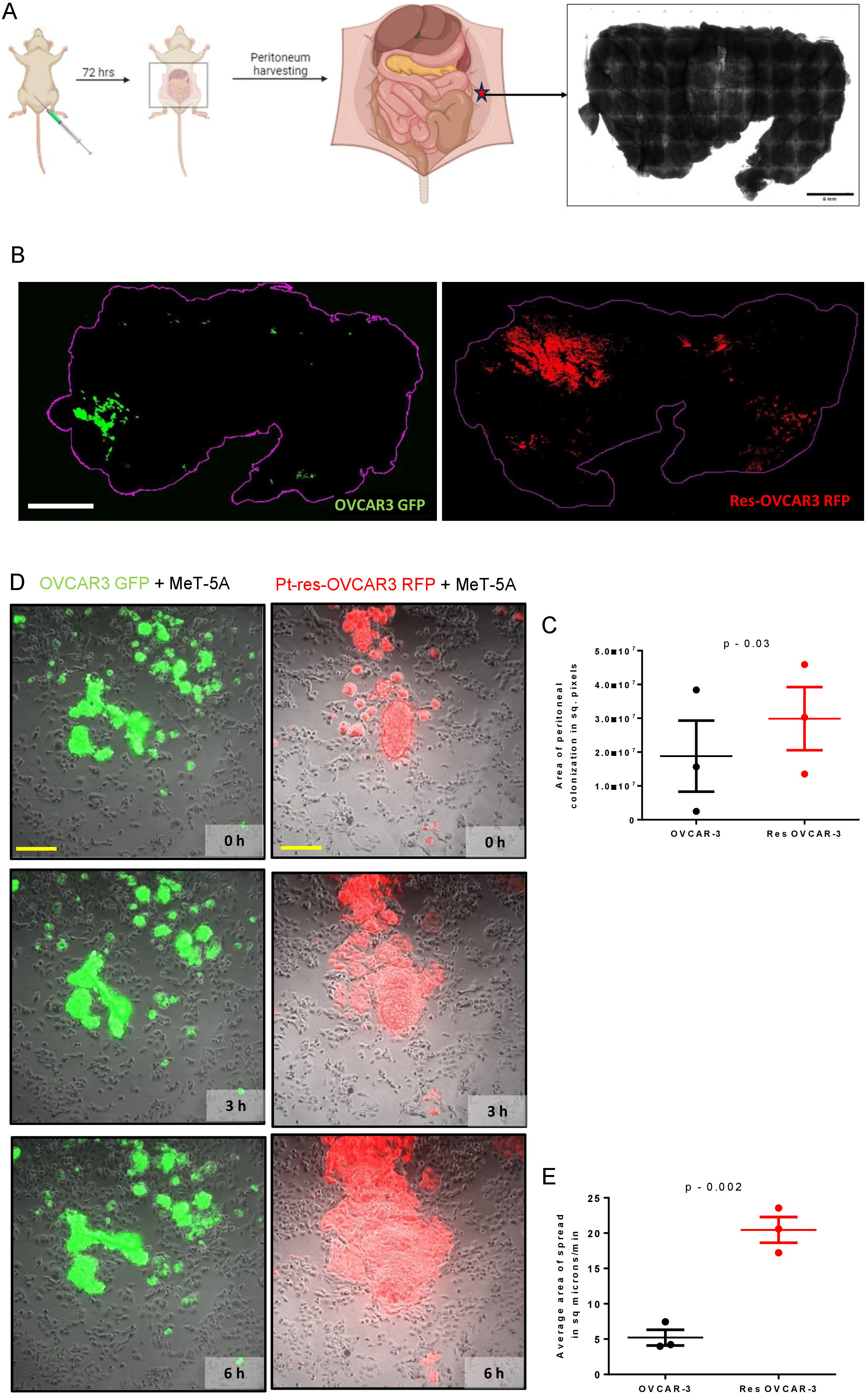
Res OVCAR-3 shows enhanced in vivo colonization and mesothelial clearance. (A) Experimental design to study peritoneal colonization. Fluorescently labeled cells (OVCAR-3 GFP, res OVCAR-3 RFP) were injected via intraperitoneal route. 72 hrs post-injection, the parietal peritoneum is harvested and imaged with tiling and stitching to identify the presence of fluorescent cells. Illustration made using BioRender. (B) Stitched fields of epifluorescent micrographs depicting the harvested parietal peritoneum (outlined in purple) showing colonization by OVCAR-3 GFP (left) and resOVCAR-3 RFP (right) cells. Scale bar - 6 mm. (C) Graph showing the area of peritoneal colonization by OVCAR-3 GFP and res OVCAR-3 RFP cells. The data represented is cumulative of 3 independent experiments, each performed with at least 2 mice per test category. Control mice were injected with PBS. Bars indicate mean ± SEM. Significance is measured using a paired t-test. (D) Photomicrographs captured using time-lapse imaging showing adhesion and spread of OVCAR-3 GFP (left) and resOVCAR-3 RFP (right) clusters on top of untransformed human coelomic mesothelial cells (MeT-5A). Scale bar – 200 μm. Refer to videos S7 and S8. (E) Graph representing the area of spread of fluorescently labeled cancer cell clusters on a monolayer of untransformed human coelomic mesothelial cells (MeT-5A) at 6 h. The data represented is cumulative of 3 independent biological experiments performed with at least 2 duplicate samples run in each set. Bars indicate mean ± SEM. Significance is measured using an unpaired student’s t-test.

We have in previous papers shown that ovarian cancer spheroids that form lumen (and are called blastuloids) tend to adhere poorly to murine peritoneal surfaces compared with when their lumen is lost due to removal of their basement membrane coats: such lumen-less ‘moruloids’ (Langthasa et al., 2021). It was, therefore, unsurprising that we observed a predominantly poor lumen formation and a moruloid phenotype in the spheroids constituted from res OVCAR-3 cells (Figure S9, photomicrographs of mature spheroids formed by OVCAR-3 and resistant OVCAR-3 variant lines). This was observed to a greater degree in the second independent line resOVCAR-3(2), which was also found to adhere poorly on Collagen I substrata compared with control OVCAR-3 (Figure S10).

### Evolved resistance and migration have distinct molecular drivers

We next sought to identify the molecular players responsible for these phenotypes, as they can prove to be valuable targets for therapeutic intervention. Here we examined the roles of CDH1 and LGALS3BP genes in developing chemo-resistance and the associated migratory properties. CDH1 is a strong epithelial marker that contributes to the epithelial properties of OVCAR-3 cells. Our results described above showed although resistant cells depart from an epithelial state to mesenchymal or amoeboid states to aid in migration, the expression of CDH1 remains higher compared to OVCAR-3 in resOVCAR-3 cells (Figure 3D). LGALS3BP is a secreted protein associated with cancer metastasis (Capone et al., 2021; Iacobelli et al., 1994) and ishighly expressed in epithelial ovarian cancer tissues compared to normal ovarian tissues (Qu et al., 2016), and early reports suggest that monitoring the serum levels of CA125 and LGALS3BP can be used to better detect ovarian cancer (Scambia et al., 1988). LGALS3BP was also found to be upregulated (1.7-fold increase) in resOVCAR-3 compared to OVCAR-3 (Figure 6A; p = 0.01 measured using an unpaired student’s t-test). Since a mechanistic understanding of the role of these markers in chemoresistance is still limited, we sought to explore their roles in the acquisition of chemoresistance and its associated emergent traits in resOVCAR-3 cells. Upon knockdown of CDH1 (Figure S11) in resOVCAR-3 cells, there was a 1.5-fold decrease in the carboplatin IC50 value of resOVCAR-3-CDH1-shRNA cells compared to the scrambled control (Figure 6B, p<0.01 significance established using unpaired student’s t-test), but there were no significant differences in the invasion properties of the CDH1 knockdown cells (Figure 6C, no significant fold difference in mean cell number per field; Figure 6D, epifluorescent micrographs of resOVCAR-3-shsc and resOVCAR-3 CDH1 KD cells stained with PI for DNA (white against black background)). The knockdown of LGALS3BP (Figure S12) in res OVCAR-3 cells showed no significant differences in carboplatin sensitivity (Figure 6E), but it decreased the invasion of resOVCAR-3-LGALS3BP-shRNA cells on collagen-I compared to the scrambled controls (Figure 6F, epifluorescent micrographs of resOVCAR-3-shsc (left) and resOVCAR-3-LGALS3BP-shRNA (right) cells stained with propidium iodide (PI) for DNA (white against black background); Figure 6G, 2.7-fold difference in mean cell number per field p=0.03 with significance calculated using a paired student’s t-test)..

**Figure 6:**
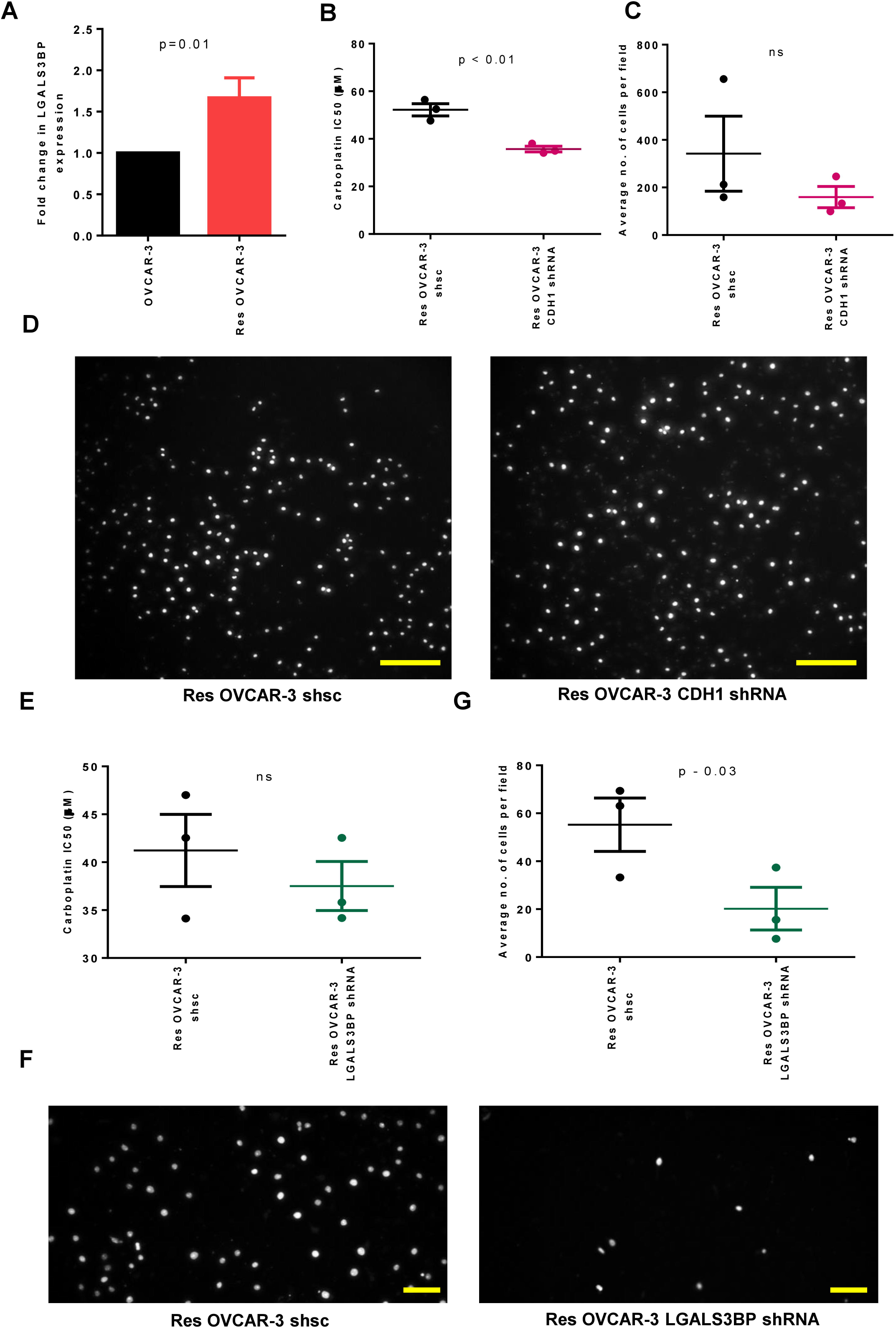
Distinct molecular players drive chemoresistance and migratory phenotypes. (A) Graph showing relative mRNA levels of LGALS3BP using qPCR in resOVCAR-3 compared to OVCAR-3 (18sRNA used as internal control). (B) Graph showing IC_50_ values for carboplatin in scrambled control (shsc) resOVCAR-3 and resOVCAR-3 CDH1 KD cells using resazurin assay. (C) Graph showing the number of resOVCAR-3 shsc and resOVCAR-3 CDH1 KD cells per field that have invaded through Collagen-I coated cell culture inserts. (D) Epifluorescent micrographs of resOVCAR-3 shsc (left) and resOVCAR-3 CDH1 shRNA (right) cells on the opposite side of the transwell stained with propidium iodide (PI) for DNA. Scale bar - 200 µm. (E) Graph showing IC_50_ values for carboplatin in scrambled control (shsc) resOVCAR-3 and resOVCAR-3 LGALS3BP KD cells using resazurin assay. (F) Epifluorescent micrographs of resOVCAR-3 shsc (left) and resOVCAR-3 LGALS3BP shRNA (right) cells on the opposite side of the transwell stained with propidium iodide (PI) for DNA. Scale bar - 100 µm. (G) Graph showing the number of control (shsc) resOVCAR-3 and resOVCAR-3 LGALS3BP KD cells per field that have invaded through Collagen-I coated cell culture inserts. Bars indicate mean ± SEM. Significance is measured using a paired t-test. (B, C, E) Bars indicate mean ± SEM. Experiments performed n ≥ 3 times. Significance is measured using an unpaired t-test.

## Discussion

The evolution of platinum resistance is the predominant therapeutic challenge in the management of EOC. Although resistance is clinically assessed through a threshold of recurrence before the six-month mark after the last dose of platinum therapy, even patients who relapse beyond the threshold eventually show poor response to platinum drugs, leading to diminishing progression-free survival (Moore et al., 2018). In this study, we chose to look at chemoresistance as one of a set of canonical histopathological tumor traits that combinedly evolve in response to selective pressures of platinum exposure. We demonstrate that such coevolution results in a more invasive phenotype, which would be responsible for the aggression of chemoresistant EOC.

A rigorous examination of the invasiveness of resistant cells in this study suggests unique morphogenetic dynamics. Given that OVCAR-3 cells are epithelioid cells with a capacity to form adherent patches, it was intriguing that the resistant cells showed even higher levels of both E-cadherin and Fibronectin. However, this is consonant with the non-monotonic variation of functional properties from epithelial to mesenchymal states; for instance, hybrid E/M cells have been reported to be the most plastic, stem-like, and metastatic (Grosse-Wilde et al., 2015; Hari et al., 2022; Pastushenko et al., 2018). Gene enrichment prediction of the bulk RNA sequencing of the carboplatin-resistant cells also pointed to the transition between these phenotypes, which is now documented in the evolution of chemoresistance across cancer types (Debaugnies et al., 2023b; Lüönd et al., 2021). However, their poor adhesion to ECM and relatively fainter basal stress fiber staining indicated the transition further extends toward an amoeboid-like state. The novelty of such a collective transitory state was evidenced by the deformable translational dynamics within smaller adherent clusters of drug-resistant cells, which also show evidence of supracellular F-actin organization at the cortex of the clusters. In contrast, within clusters of sensitive ovarian cancer cells, F-actin is localized subcellularly to cell cortices. These observations suggest that clusters of resistant cells are organized like a single giant cell that jiggles and moves en masse. Such behavior has been observed for cell clusters in non-adhesive environments and has been referred to as collective amoeboid migration (Pagès et al., 2022). We observe the collective amoeboid migration within suspended spheroids of resistant cancer cells, which showed mesoscale-level pseudopod-like extensions and retractions while translating across polymerized fibrillar Collagen I surfaces.

Collective low-adhesion translation of multicellular clusters has been elegantly studied by Beaune and coworkers, who used a combination of particle imaging velocimetry and traction force microscopic measurements to delineate how such clusters translate on low stiffness substrate but dismantle and spread on rigid surfaces (Beaune et al., 2018). The velocity of such clusters is driven by the difference in spreading parameters at the receding and moving edge of the clusters, which in turn depends on the traction exerted by these clusters on the surface, leading to an irreversible modification of the surface at the rear end. However, in their time lapses, they do not capture the formation of pseudopodia-like projections, nor drastic deformation in cluster shape of the nature we observe: this sheds light on the cell-contextual nature of collective behavior. The movement of our clusters may be driven by asymmetricity in both intercell adhesion strengths due to increased E-cadherin levels and cohesiveness due to fibronectin. We look forward to testing this hypothesis in future studies using traction force microscopic measurements in spheroids from cells expressing live fibronectin and E-cadherin reporters.

Cells can deploy amoeboid migration to move faster than when they are mesenchymal. Such features help the cell to adapt and migrate through narrow spaces in the ECM or even exert a force sufficient to deform the ECM (Driscoll et al., 2024; Jeong et al., 2012; McGrail et al., 2014; Śliwa et al., 2024). High levels of the Rho/ROCK signaling pathway are associated with and promote amoeboid migration (Graziani et al., 2022; Sanz-Moreno et al., 2008). In ovarian cancer, the overexpression of Rho/Rock kinases increases cancer cell dissemination and increases their resistance to cisplatin, alluding to the potential of kinase-specific inhibitors as having therapeutic potential (Horiuchi et al., 2008; Ohta et al., 2012; Śliwa et al., 2024). In fact, melanoma cells that are resistant to tyrosine kinase inhibitors show higher levels of myosin II, which is associated with the amoeboid migratory phenotype (Orgaz et al., 2020). The actomyosin-driven deformability allows such cells to metastasize faster. This is consistent with our observation that resistant cells colonize and spread further on peritoneal surfaces. Moreover, amoeboid cells express markers that strongly overlap with those typical of epithelial-to-mesenchymal transitions, such as RHO-ROCK signaling and TGF-β-SMAD (Cantelli et al., 2015). The increase in mesenchymal marker fibronectin in our resistant cells and the co-existence of lower cell-substrate adhesion with higher gene enrichment-based scores for EMT and EMAT is consistent with the amoeboid state being related to the epithelial-mesenchymal spectrum. Even so, we propose that the resistant cell migratory dynamics represent a unique transitory state due to its upregulation of E-cadherin (downregulated in classical amoeboid and mesenchymal cells), contributing to its collective nature.

A careful study of the transitory dynamics will be the focus of subsequent investigations. Resistant cell populations may represent heterogeneous mixtures of co-existent cell populations with distinct cell states. On the other hand, we cannot rule out plasticity among different cell-states where the transition rates may depend on cell confluence, local rheological heterogeneity, and other ecological interactions among diverse phenotypes (Gasior et al., 2019; Jain et al., 2024; McFaline-Figueroa et al., 2019). We aim to interrogate these questions using live expression markers of cell states as well as undertake single-cell RNA sequencing and multiplexed immunocytochemistry to dissect subpopulation level behaviors. The factors governing the rates of these cell-state transitions would help us better understand and eventually control co-evolution trajectories in a heterogeneous population subjected to therapeutic stress.

The temporal scale over which the divergence in the phenotype of the resistant from sensitive cancer cells took place was several weeks: it is, therefore, conceivable that distinct molecular mechanisms drove resistance to the selective pressures of carboplatin exposure, and other traits may have evolved subsequently through consequential ‘passenger’ changes. This led us to investigate the effects of E-Cadherin, and a known marker of mesenchymal state associated with high-grade serous ovarian cancer LGALS3BP, on the traits of drug resistance and invasion. The association of E-Cadherin depletion with loss of drug resistance indicates it plays an early and causal role in the acquisition of resistance to carboplatin. In contrast, the association of LGALS3BP levels with invasion but not carboplatin resistance suggests its elevation during resistance evolution may be temporally distal to the elevation in levels of E-cadherin. Future studies will probe the correlations or lack thereof in the temporal evolution of individual traits and their molecular mediators.

Finally, the observations reported in this study are based on selection experiments on HGSOC-mimicking lines under cultivation as monolayers. While the results from the system are strongly resonant at the cellular and molecular level with existing literature, a better understanding of the chemoresistance mechanisms can be gained by developing patient-derived xenograft (PDX) models of chemo-sensitive and chemoresistant tumors. The validation of such novel migratory dynamics within tumor microenvironment-incorporating experimental models should result in the identification of unique therapeutic targets for the refractory form of cancer.

## Materials and Methods

### Cell culture

The human ovarian cancer cell line OVCAR-3 was a kind gift from Professor Rajan R. Dighe, Indian Institute of Science, and COV362 was purchased from SIGMA (cat no. 07071910). The human immortalized untransformed mesothelial cell line MeT-5A was purchased from ATCC (CRL-9444). OVCAR-3 and res OVCAR-3 cells were maintained in RPMI-1640 medium (AL162A; HiMedia) supplemented with 20% FBS (10270; Gibco), while the COV362 and res COV362 cells were cultured in DMEM (AL007A; HiMedia) supplemented with 2 mM glutamine and 10% FBS. MeT-5A cells were maintained in M199 medium (AL014A; HiMedia) supplemented with 10% FBS, 5.053 µg/ml insulin, 0.5 ug/ml hydrocortisone, 2.6 ng/ml sodium selenite, 10 ng/ml human epidermal growth factor (hEGF), 20 mM HEPES. Cells were subcultured in a 1:4 ratio at approximately 80% confluence using 0.05% trypsin-EDTA (TCL007; HiMedia) solution for trypsinization and incubated in a humidified incubator with 5% carbon dioxide at 37 °C.

### Establishing resistant cell lines

OVCAR-3 and COV362 cells at 70-80% confluence were treated with IC_50_ dose carboplatin (Cayman Chemicals - 13112) for 2 hours, after which the drug was removed, and cells were allowed to recover. Once the cells attained 80% confluence after recovery, they were trypsinized and re-plated at a 1:4 ratio. The treatment was repeated at 70-80% confluence, followed by recovery, and the cycle was repeated till the desired level of chemoresistance was established. Acquisition of chemoresistance was ascertained by assessing viability in the presence of carboplatin treatment using resazurin assay.

### Resazurin assay

The viability of cells treated with chemotherapeutic agents was ascertained using a cell-permeable redox indicator resazurin (Sigma-Aldrich - R7017). Resazurin, a blue non-fluorescent dye, is reduced to a fluorescent compound resorufin by metabolically active viable cells at rates proportional to the number of viable cells present. Cells were seeded in 96 well plates at a density of 3000 cells/well and, after 12 hrs, treated with desired concentrations of the chemotherapeutic agent for 72 hrs. 1 µg/mL puromycin (Sigma-Aldrich – P8833) treatment was used as a positive control for cell death. After incubation, resazurin was added to each well at a final concentration of 10 µg/µL and further incubated for 1.5 hrs at 37°C. Resorufin fluorescence was quantified using a microplate fluorometer (Tecan Infinite M Plex) with a 560 nm excitation/590 nm emission filter set. Prism software (GraphPad Prism 6.0) was used to determine IC50 values.

### Proliferation assay

OVCAR-3 and resOVCAR-3 cells were seeded at a density of 30,000 cells/well of a 12-well plate. After 12 hrs, cells were treated with 30 µM carboplatin and incubated for 72 hours at 37°C. After incubation, the cells were trypsinized and counted using a hemocytometer, and fold change in the total number of viable cells was calculated.

### Collagen-I polymerization

Rat tail collagen (A10483-01; Gibco) neutralization was performed on ice by adding 10X DMEM and 0.1 N NaOH in a ratio of 333:100:5 (Collagen: 10X DMEM: 2N NaOH). The neutralized collagen was diluted with 1X PBS or medium to obtain the required collagen concentration.

### Transwell invasion assay

Polycarbonate transwell inserts with 8 μm pore-size (HiMedia; TCP083) were coated with 200 μg/mL neutralized rat tail collagen. Cells were seeded inside each transwell in 200 μL of serum-free medium. The number of cells seeded varied according to the cell line (OVCAR-3 – 30,000 cells/transwell, COV362 – 10,000 cells/transwell). The bottom well or the reservoir was filled with 1 mL of serum-containing medium and incubated for 9 hours (COV362 cells) or 24 hours (OVCAR-3 cells) at 37°C. The medium from the transwells was carefully removed and washed with 1X PBS, followed by fixing the cells using 100% methanol for 10 minutes at −20 °C. The fixed cells were rewashed with 1X PBS, and non-invading cells and the collagen matrix were carefully removed with moistened cotton Q-tips. The transwells were stained with 50 μg/mL of propidium iodide for 5-10 minutes and washed to remove excess dye. The membranes were imaged under a fluorescence microscope using a 10X magnification objective, and the number of cells in each field was quantified.

### Wound Healing / Scratch Assay

50,000 cells were seeded per well in an 8-well-chamber plate and incubated at 37°C for 24-48 hours till the cells formed a confluent monolayer. A scratch was made on the surface of the confluent layer of cells using a 2 μL pipette tip. The empty patch was observed for 24 hours using a time-lapse videography setup equipped with a 5% CO2 incubator, and the rate at which cells repopulate the patch was determined.

### Spheroid culture

Tissue culture dishes were coated with 3% poly-2-hydroxyethyl methacrylate (poly HEMA; P3932, Sigma-Aldrich) solution prepared in 95 % absolute ethanol and allowed to polymerize under sterile conditions at room temperature overnight. The dishes were then sterilized under ultraviolet radiation overnight. Cells were seeded in a defined medium consisting of the base medium used for the cell line supplemented with 0.5 μg/ml hydrocortisone (Sigma-Aldrich, 33 H0888), 250 ng/mL insulin (Sigma-Aldrich, I6634), 2.6 ng/mL sodium selenite (Sigma-Aldrich, S5261), 27.3 pg/mL estradiol (Sigma-Aldrich, E2758), 5 μg/mL prolactin (Sigma-Aldrich L6520, 10 μg/mL transferrin (Sigma-Aldrich, T3309). The cultures were maintained in a 5% CO2 incubator at 37°C for a minimum of 24 hrs or longer as per the requirements of the experiment.

### Adhesion assay

96 well plates were coated with 50 μg/mL neutralized rat tail collagen. Cells were seeded at 10,000 cells/well density and incubated at 37°C for 30 minutes. The wells were thoroughly washed 5-6 times using 1X PBS to remove floating cells. The adherent cells were fixed using 100% methanol at −20 °C for 10 minutes. Following 1-2 1X PBS washes, the fixed cells were stained with 50 μg/mL of propidium iodide for 10 minutes. After staining, the cells were rewashed with 1X PBS, and the fluorescence intensity of the adhered cells was recorded using a microplate fluorometer.

### Immunostaining

Eight well-chamber plates containing cells or cells cultured on ECM were fixed using 3.7% formaldehyde (24005; Thermo Fisher Scientific) at 4°C for 20 min. Fixed cells were permeabilized using 0.5% Triton X-100 (MB031; HiMedia) for 1–2 hr at RT to enable effective entry and uniform exposure to the antibodies. Permeabilization was followed by a blocking step using 1X PBS with 0.1% Triton X-100 and BSA (MB083; HiMedia) for 45 min at RT. The cells were then incubated with the primary antibody (diluted according to the manufacturer’s protocols) overnight at 4°C. This was followed by three washes with 0.1% TritonX-100 in 1XPBS (5 minutes each). Secondary antibody incubation along with phalloidin [Alexa Fluor™ 488 Phalloidin (A12379, Thermo Fischer Scientific), Alexa Fluor™ 568 Phalloidin (A12380, Thermo Fisher Scientific)] was performed at RT for 2 hr under dark conditions. Cells were counterstained with DAPI (1:1000 dilution) (D1306; Thermo Fisher Scientific) and washed away after 15 min. Subsequent processes, including washes, were carried out in the dark. Images were captured using an Olympus IX73 microscope with an Aurox spinning disk confocal setup or an Andor Dragonfly (CR-DFLY-502) confocal imaging platform. Images were processed using ImageJ software. The antibodies used in our studies are against FN1 (E5H6X, Cell Signaling Technology, Inc.) and CDH1 (24E10, Cell Signaling Technology, Inc.). Each experiment performed included negative controls, which were devoid of primary antibodies.

### In vivo peritoneal colonization study

GFP/RFP labeled OVCAR-3 and res OVCAR-3 cells were used for this assay. 200 μL cell suspension containing 7×10^6^ – 10×10^6^ cancer cells were injected unilaterally into the peritoneal cavity of 6-8 weeks old female athymic nude mice. Mice subjected to IP injections of PBS were used as controls. 72 hours post-injection, mice were euthanized by cervical dislocation. The abdomen was surgically cut open, and the outer skin was teased apart from the peritoneal lining. The peritoneal membrane was harvested and placed stretched-out on 60 mm dishes. Special care must be taken to ensure that the side of the membrane facing the organs faces the bottom of the dish. The membrane was kept in 1X PBS solution and imaged as live tissue without fixation. The entire harvested peritoneal membrane was imaged in tiling mode in an epifluorescence microscope (Olympus IX83). The animal studies described here were carried out according to the guidelines of the institutional review board and in agreement with the ethical guidelines of IISc (IAEC approved protocol No.: CAF/ethics/635/2018).

### Co-culture study

Non-fluorescent MeT-5A cells were seeded in eight-well chambered cover glasses at 40,000 cells/well density and incubated for 24 hours at 37 °C to form 60-70% confluent monolayers. Simultaneously, fluorescently labeled cancer cells were seeded in poly-HEMA-coated dishes to form spheroids. After 24 hours, the fluorescent OVCAR-3 or resOVCAR-3 spheroids were added to the Met-5A monolayers. Time-lapse videography setup equipped with a 5% CO2 incubator was used to observe the interaction between the ovarian cancer cells and the Met-5A cells. The acquired images were processed using ImageJ to analyze the area of spread of cancer cell clusters.

### Spheroid adhesion assay

OVCAR-3 and resOVCAR-3 cells were cultured in poly-HEMA coated dishes for seven days to form lumen-containing spheroids. Spent media was replaced every two days. Mature spheroids were seeded on top of neutralized 1 mg/mL collagen-I in an 8-well-chamber plate. After counting the initial number of clusters present, the plate was incubated for 5 hours, after which the collagen gels were washed with 1X PBS, and the remaining number of spheroids were counted to calculate the percentage of cells that have adhered to the collagen-I gel.

### Time-lapse imaging

Cell culture setups specific to the experiment were prepared in eight-well-chambered cover glasses placed in the Tokai stage-top incubator of an Olympus IX73 microscope. The stage-top incubator was maintained at 37 °C, with 5 % CO2, and sterile water was added to the designated humidification space. The CellSens imaging software was used to control all the connected parts associated with imaging, including the Orca Flash LT plus camera (Hamamatsu). Acquisition and exposure settings were made according to the channels used, and the multipoint time-lapse imaging was carried out for 24-48 hours, with a time interval of 10-30 minutes.

### Gene expression profiling

Cells were lysed using a Trizol reagent (Takara, Japan). RNA was isolated by the chloroform-isopropanol extraction method. All reagents were molecular grade and purchased from Merck. Per the manufacturer’s protocol, 1 μg of total RNA was reverse transcribed using the Verso™ cDNA synthesis kit (Thermo Scientific, AB-1453). Real-time PCR was performed with 1:10 diluted cDNA using the SYBR green detection system (Thermo Fischer Scientific, F415L) and Rotorgene Q (Qiagen, 9001560) using the 18S rRNA gene as the internal control. Relative gene expression was calculated using the comparative Ct method. Appropriate no template and no RT control were included in each experiment. All the samples were analyzed in triplicates and repeated three times independently. Primer sequences of CDH1, FN1, LGALS3BP, and 18s rRNA genes used in this study are as follows: 18S forward - GTAACCCGTTGAACCCCATT, 18S reverse - CCATCCAATCGGTAGTAGCG, CDH1 forward - ACCACCTCCACAGCCACCGT, CDH1 reverse - GCCCACGCCAAAGTCCTCGG, FN1 forward - CAAGCCAGATGTCAGAAGC, FN1 reverse - GGATGGTGCATCAATGGCA, LGALS3BP forward - AGGTACTTCTACTCCCGAAGGA, LGALS3BP reverse - GGCCACTGCATAGGCATACA.

### Genetic perturbation of CDH1 and LGALS3BP genes

The CDH1 and LGALS3BP shRNA clones were obtained from the MISSION shRNA library (Sigma Merck). The plasmids containing shRNA or scrambled controls were packaged into lentiviruses using pMD2.G and psPAX2 packaging vectors. The packaging vectors were a kind gift from Professor Deepak K Saini, DBG, Indian Institute of Science. The plasmids were transfected into 293FT cells (R70007; Thermo Fisher Scientific) using TurboFect (R0533; Thermo Fisher Scientific) in the following ratio - 2000 ng psPAX2: 500 ng pMD2.G: 2000 ng shRNA plasmid. 293FT cells were cultured in DMEM supplemented with 10% FBS. The conditioned medium of the transfected cells containing viral particles was collected at 48 and 72 hrs and filtered through a 0.2 μm filter followed by a concentration process using the Lenti-X concentrator according to the manufacturer’s protocol (631232; TaKaRa). The concentrated virus was stored as aliquots at −80°C until use. resOVCAR3 cells were seeded in a 6-well plate at 50−60% confluence and transduced with viral particles containing shRNA or scrambled control and polybrene (8 μg/ml). Transduced cells were selected using 5 μg/ml puromycin after 72 hrs. The knockdown of the gene was confirmed using real-time qPCR.

### RNA-Seq Data Analysis

RNA-Seq data was processed using the nf-core RNA-Seq pipeline (version v3.14.0) (Ewels et al., 2020), a standardized and automated pipeline designed for comprehensive RNA-Seq data analysis. The bam files obtained from the nf-core RNA-Seq pipeline were then used to generate an expression matrix using featureCounts v2.0.1. (Liao et al., 2014). CPM counts for plotting were generated using EdgeR after TMM normalization (Robinson et al., 2010). PyDESeq2 (Muzellec et al., 2023) was used for differential gene expression. A cut-off of Log2FC≥ 1 and fdr cut-off of 0.05 was used for selecting 1457 upregulated genes, and Log2FC≤ −1 and fdr cut-off of 0.05 was used for selecting 1376 downregulated genes. All the results were plotted using Python.

### single sample Gene Set Enrichment Analysis (ssGSEA)

The scoring of the samples was done using the GSEApy implementation (Fang et al., 2023) of the single sample gene set enrichment analysis (ssGSEA) algorithm (Barbie et al., 2009). The genes are ranked based on the transcript per million (TPM) levels, and the enrichment scores are calculated as the area between the empirical cumulative distribution functions (ECDFs) of the genes in the geneset and the background genes. The normalized enrichment scores were used for further analysis. The signature for epithelial, mesenchymal, partial-epithelial-mesenchymal transition (pEMT), and epithelial-mesenchymal-amoeboid transition (EMAT) signatures were obtained from previous reports (Emad et al., 2020; Puram et al., 2017; Tan et al., 2014). The p-values were obtained using two-sided Welch’s t-test (independent samples with unequal population variance).

### Gene Ontology analysis

Gene ontology analysis was performed using the ShinyGO v0.80 (Ge et al., 2020) platform for gene enrichment analysis, using the default settings for p-value cut-off after selecting human species.

### Statistical analysis

All experiments were performed with a minimum of two technical replicates and were independently repeated at least thrice. GraphPad Prism 6.0 was used for data analysis and representation. Results are represented as mean ± SEM unless stated otherwise. Statistical significance was estimated using an unpaired t-test with Welch’s correction in most cases unless mentioned otherwise.

## Acknowledgments

We would like to thank Satyarthi Mishra and Kottpalli Vidhipriya (IISc) for their valuable help in optimizing the co-culture and wound healing assays respectively. We would also like to thank the Central Animal Facility (CAF), IISc for help with the animal experiments. This work was supported by the India Alliance DBT Wellcome Trust Fellowship (IA/I/17/2/503312) awarded to RB. It was also supported by the John Templeton Foundation (no. 62220), the Indo-French Centre for the Promotion of Advanced Research (CEFIPRA grant: 69T08-2), and the International Emerging Actions (328003) to RB. SG and AG acknowledge their fellowship support from the Ministry of Education, Government of India. MKJ was supported by Param Hansa Philanthropies. HBV was supported by the Prime Minister’s Research Fellowship (PMRF) awarded by the Government of India. RS and NV acknowledge intramural funding support from NCBS-TIFR. The opinions expressed in this paper are those of the authors and not those of the John Templeton Foundation.

## Conflict of Interest Statement

The authors declare no competing interests.

## Notes

### Competing Interest Statement

The authors have declared no competing interest.

