## Supplementary figures for "Collective amoeboid dynamics drives colonization of drug-resistant ovarian cancer cells"

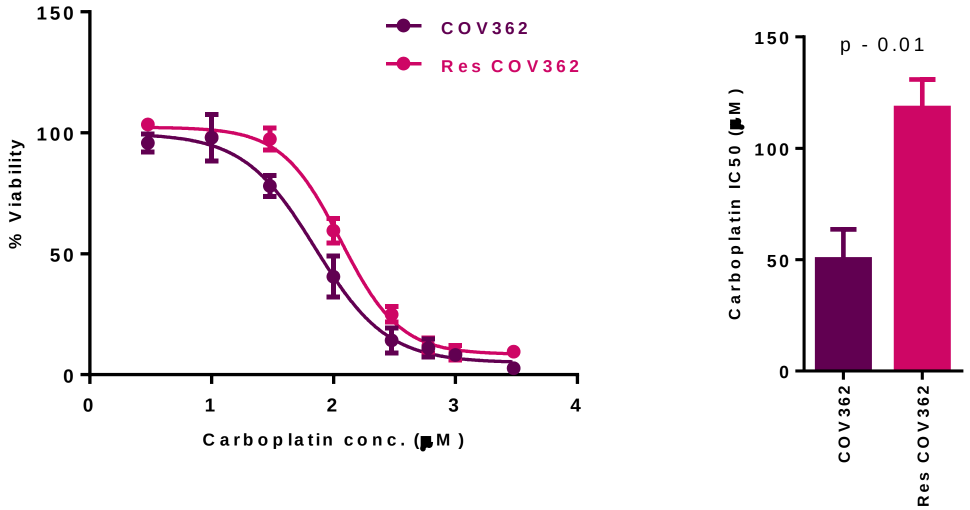


Figure S1: Graphs showing the viability of COV362 and resCOV362 at different concentrations of carboplatin treatment evaluated using resazurin assay. Each point in the viability curves and bars indicates mean ± SEM, n=4. Significance was measured by an unpaired student's t-test.

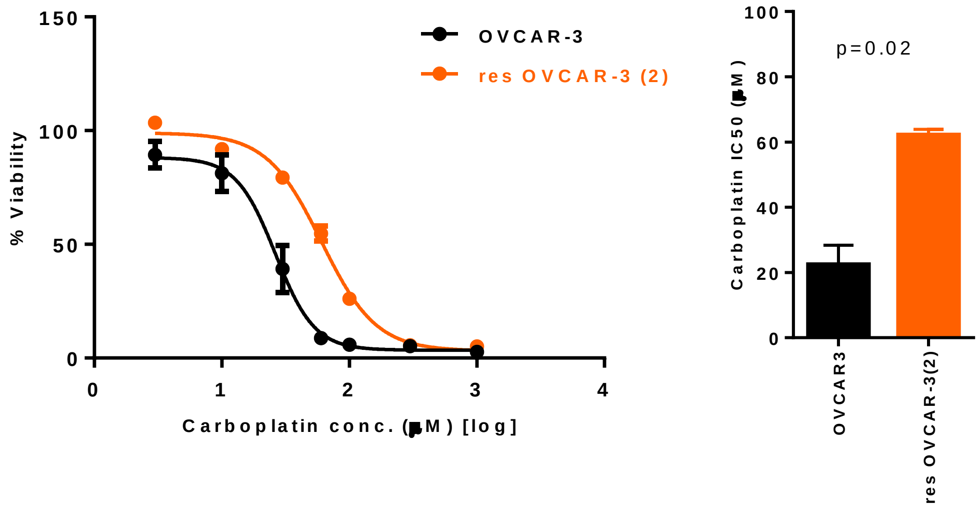


Figure S2: Graphs showing the viability of OVCAR-3 and resOVCAR-3(2) at different concentrations of carboplatin treatment evaluated using resazurin assay. Each point in the viability curves and bars indicates mean ± SEM, n=2. Significance was measured by an unpaired student's t-test.

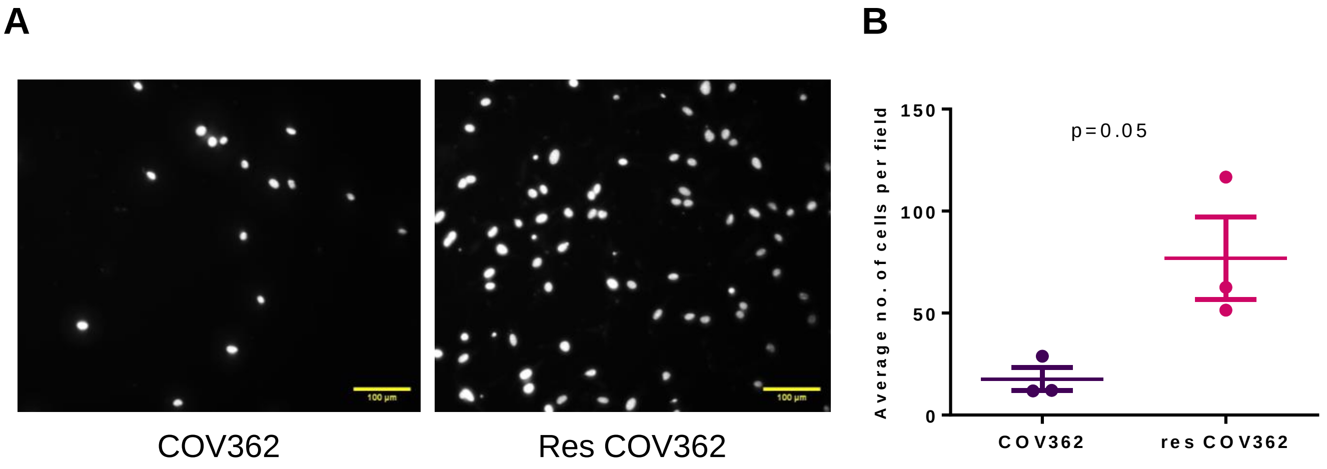


Figure S3: Invasion of COV362 and resCOV362 cells through Collagen-I coated cell culture inserts. (A) Epifluorescent micrographs of COV362 (left) and resCOV362 (right) cells stained with PI for DNA on the lower side of the transwell insert. Scale bar - 100 µm. (B) Graph showing the number of cells invaded per field. n = 3, bars represent mean ± SEM. Significance was measured by an unpaired student's t-test.

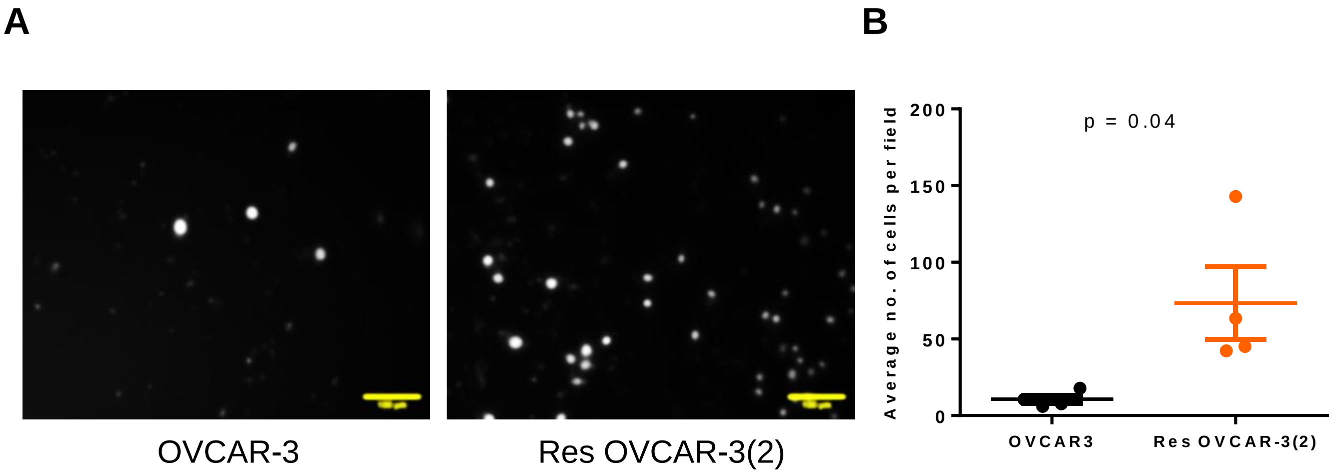


Figure S4: Invasion of OVCAR-3 and resOVCAR-3(2) cells through Collagen-I coated cell culture inserts. (A) Epifluorescent micrographs of OVCAR-3 (left) and resOVCAR-3(2) (right) cells stained with PI for DNA on the lower side of the transwell insert. Scale bar - 100 µm. (B) Graph showing the number of cells invaded per field. n = 4, bars represent mean ± SEM. Significance was measured by an unpaired student's t-test.

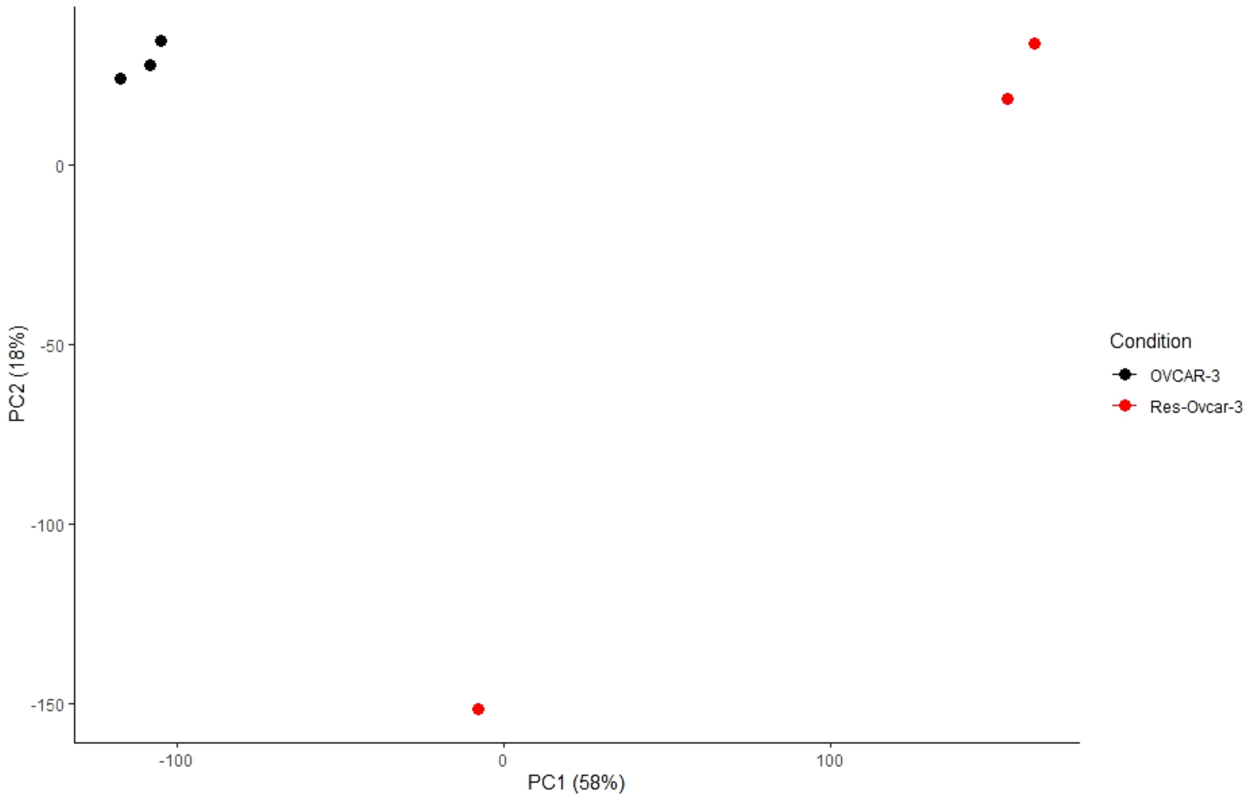


Figure S5: Principal component analysis plot showing the proximity of overall cell states of the replicates of OVCAR-3 and resOVCAR-3.

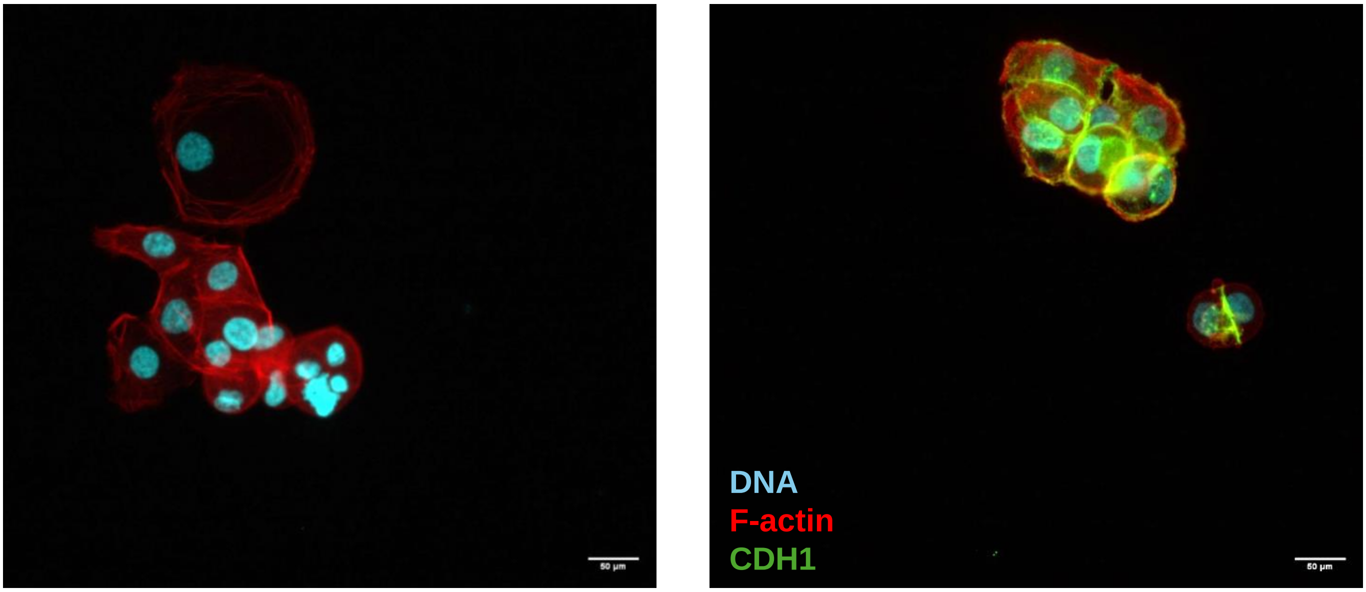


No primary antibody control

CDH1 stained

Figure S6: Fluorescent confocal micrographs of resOVCAR-3 cells cultured as monolayers stained for E-cadherin (right) and no primary antibody control (left) (green), counterstained with F-actin (phalloidin; red) and DNA (DAPI; cyan). Experiments performed n $\geq$ 3 times. Scale bar – 50 µm

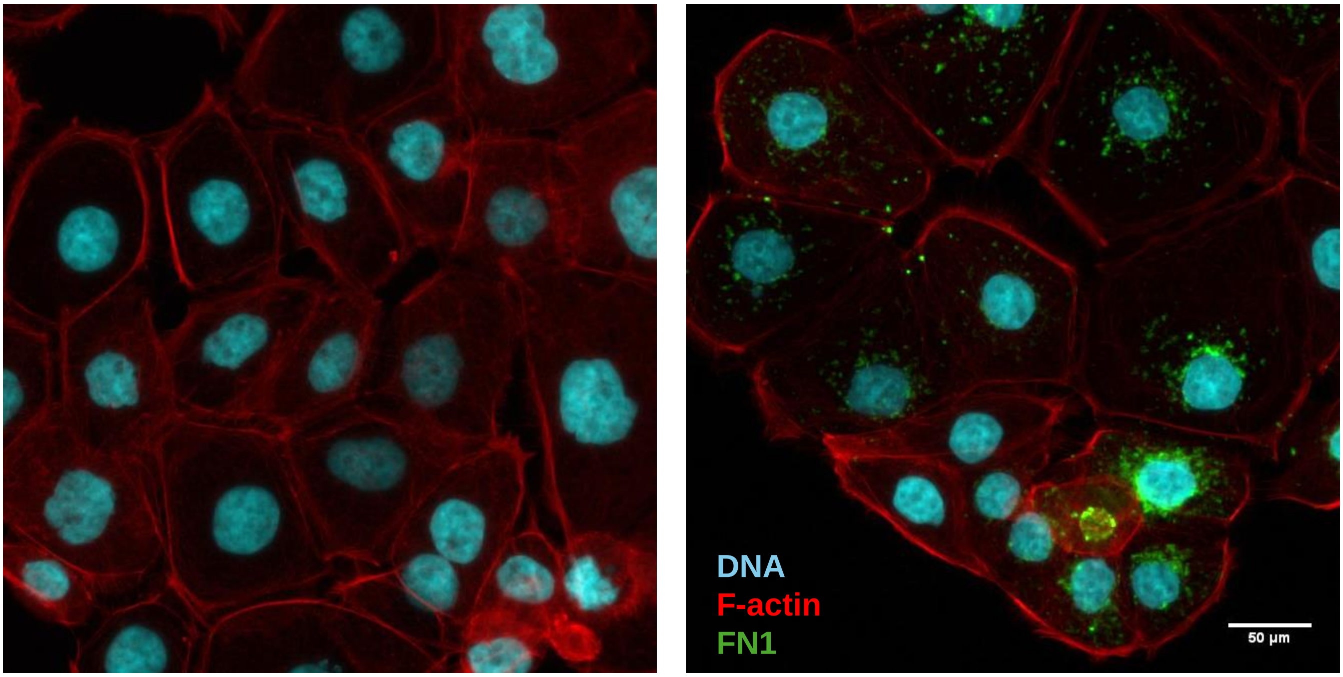


No primary antibody control

FN1 stained

Figure S7: Fluorescent confocal micrographs of resOVCAR-3 cells cultured as monolayers stained for fibronectin (right) and no primary antibody control (left) (green), counterstained with F-actin (phalloidin; red) and DNA (DAPI; cyan). Experiments performed n $\geq$ 3 times. Scale bar – 50 µm . Refer to supplementary controls

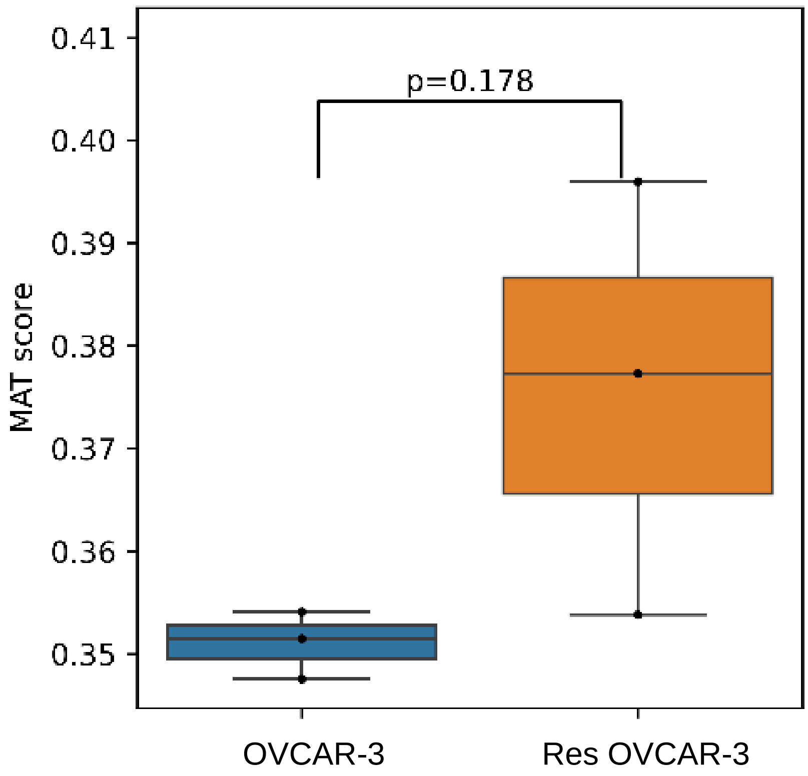


Figure S8: ssGSEA analysis plot from the RNA sequencing data presented in Figure 2 indicating the expression of curated MAT markers in OVCAR-3 and resOVCAR-3 cells.

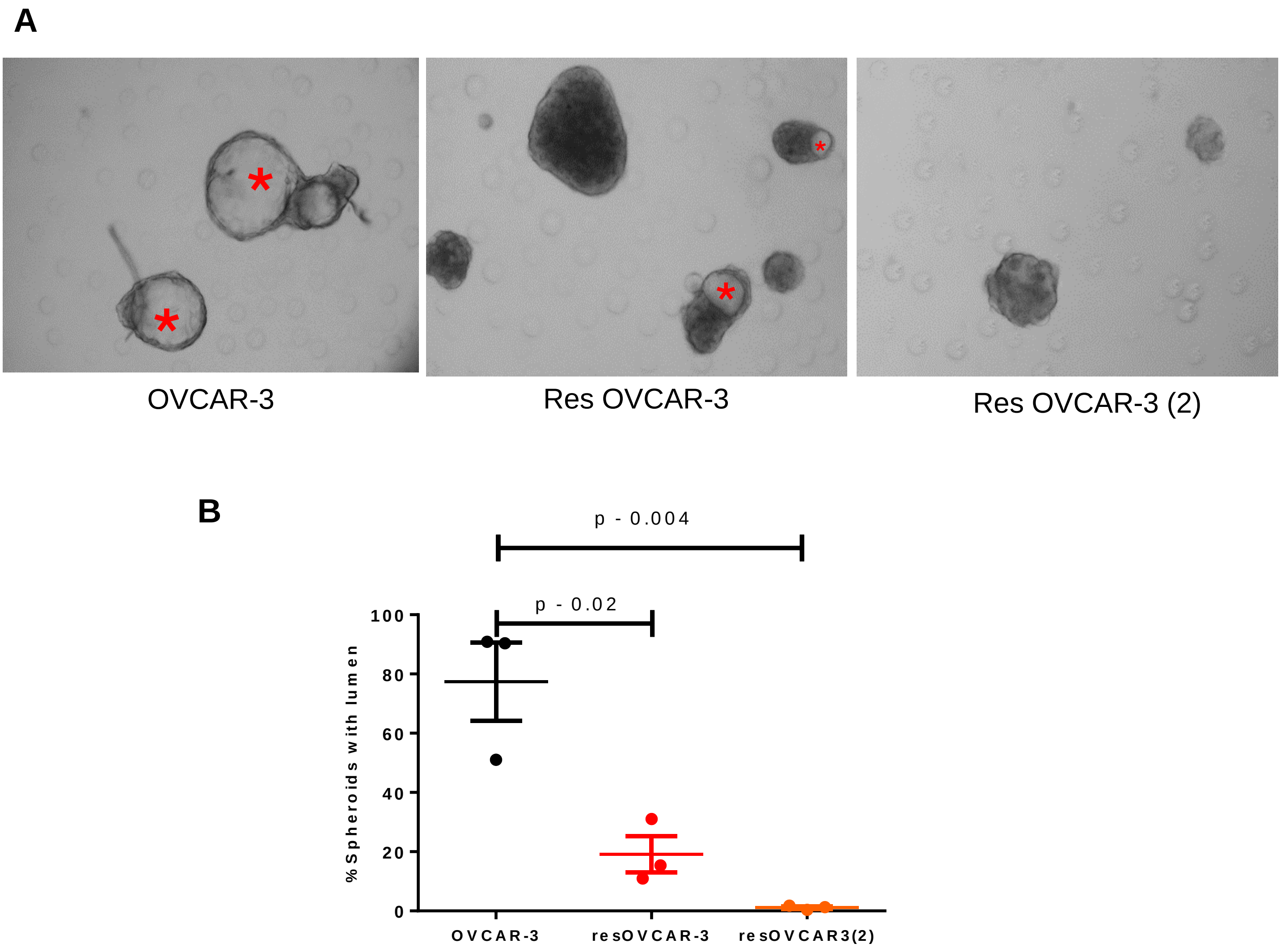


Figure S9: (A) Photomicrographs of mature spheroids formed by OVCAR-3 (left), resOVCAR-3 (middle), and resOVCAR-3 (2) cells. The asterisk indicates the lumen. (B) Graph showing the percentage of mature spheroids that form lumen. n=3, bars indicate mean ± SEM. Significance is measured using an unpaired student's t-test.

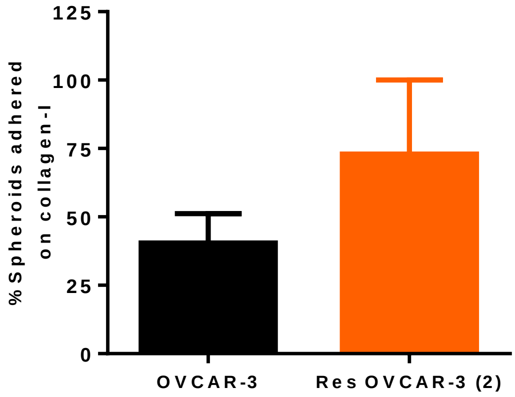


Figure S10: Graph showing adhesion of mature OVCAR-3 and resOVCAR-3(2) spheroids on 1 mg/mL collagen scaffolds at 5 h. n=2. Bars indicate mean ± SEM.

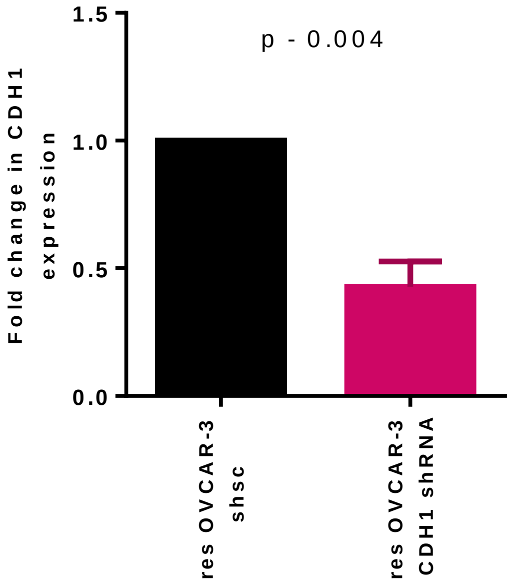


Figure S11: Graph showing decreased mRNA levels of CDH1 in resOVCAR-3-CDH1-shRNA compared to resOVCAR-3-shsc (18sRNA used as internal control) observed using qPCR analysis. Experiments performed n = 3 times. Bars indicate mean ± SEM. Significance is measured using an unpaired t-test.

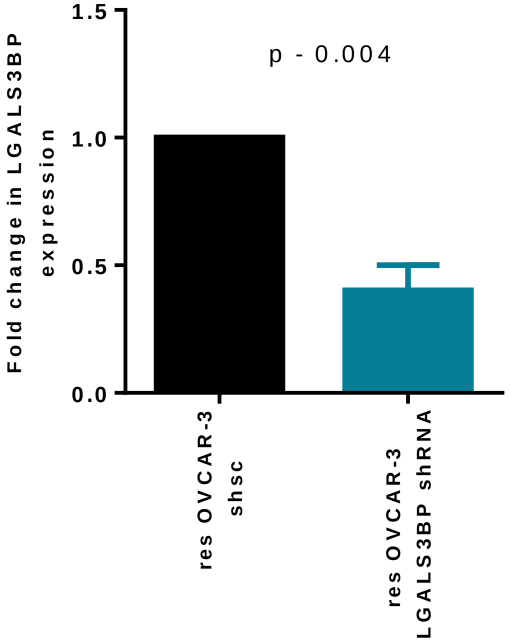


Figure S12: Graph showing decreased mRNA levels of LGALS3BP in resOVCAR-3-LGALS3BP-shRNA compared to resOVCAR-3-shsc (18sRNA used as internal control) observed using qPCR analysis. Experiments performed n = 3 times. Bars indicate mean ± SEM. Significance is measured using an unpaired t-test.

| Gene | $log_2FC$ | adj p value |
|---|---|---|
| IGF1 | -4.29384 | NaN |
| IGF2 | -1.71793 | 3.68E-05 |
| ITGB8 | -1.07129 | 5.07E-03 |
| COL1A2 | -0.80198 | 2.85E-03 |
| SPP1 | -0.72047 | 1.14E-03 |
| TNC | -0.59335 | 3.11E-09 |
| FGFR2 | -0.463 | 1.50E-04 |
| ITGB5 | -0.09236 | 5.13E-01 |
| COL1A1 | 0.025057 | 9.18E-01 |
| LAMA5 | 0.232369 | 6.35E-02 |
| LAMC1 | 0.263199 | 1.26E-02 |
| ITGB1 | 0.39475 | 5.50E-06 |
| COL4A2 | 0.419552 | 7.57E-07 |
| EFNA1 | 0.554422 | 1.30E-09 |
| EPHA2 | 0.753812 | 5.54E-20 |
| LAMC2 | 0.759742 | 5.64E-18 |
| ITGA6 | 1.024654 | 4.20E-19 |
| ITGA3 | 1.077559 | 2.01E-41 |
| ITGB4 | 1.788586 | 4.62E-72 |
| FN1 | 2.002223 | 4.89E-87 |

Table S1: Table depicting the $log_2$fold change and adjusted p values of 20 genes related to the PI3K pathways (Carvalho et al., 2022) in resOVCAR-3 compared to OVCAR-3, obtained using DESeq2.

| Gene | $log_2FC$ | adj p value |
|---|---|---|
| BCL6 | -1.84718 | 7.94E-42 |
| PPARGC1A | -1.09944 | 5.34E-02 |
| PMS2 | -0.59251 | 1.96E-09 |
| MLH1 | -0.42227 | 9.39E-05 |
| ATP7A | -0.40775 | 1.05E-01 |
| OGT | -0.2381 | 4.37E-02 |
| MSH6 | -0.1027 | 2.92E-01 |
| USP14 | 0.141667 | 1.76E-01 |
| ERCC1 | 0.35793 | 1.30E-03 |
| ATP7B | 0.534748 | 3.80E-03 |
| FBN1 | 0.787349 | 2.59E-10 |

Table S2: Table depicting the $log_2$fold change and adjusted p values of a curated set of 10 genes chosen based on their dysregulation in drug-resistant ovarian cancer progression (Ortiz et al., 2022) in resOVCAR-3 compared to OVCAR-3, obtained using DESeq2.

Video S1: Time-lapse video showing scratch disappearance in OVCAR-3 cell layers. Total video duration – 18 hrs. Experiments performed n≥3.

Video S2: Time-lapse video showing scratch disappearance in resOVCAR-3 cell layers. Total video duration – 18 hrs. Experiments performed n≥3.

Video S3: Time-lapse video showing bright field imaging of 24-hr old OVCAR-3 spheroids adhering and growing on top of 1 mg/mL Collagen-I scaffolds. Total video duration – 24 hrs (HH:MM). Experiments performed n ≥ 3 times.

Video S4: Time-lapse video showing bright field imaging of 24-hr old resOVCAR-3 spheroids adhering and growing on top of 1 mg/mL Collagen-I scaffolds. Total video duration – 24 hrs (HH:MM). Experiments performed n ≥ 3 times.

Video S5: Time-lapse video showing fluorescence imaging of GFP expressing OVCAR-3 cells. Total video duration – 40 hrs (HH:MM). Experiments performed n ≥ 3 times.

Video S6: Time-lapse video showing fluorescence imaging of RFP expressing resOVCAR-3 cells. Total video duration – 40 hrs (HH:MM). Experiments performed n ≥ 3 times.

Video S7: Time-lapse video showing adhesion and spread of OVCAR-3 GFP clusters on top of untransformed human coelomic mesothelial cells (MeT-5A). Total video duration – 44 hrs (HH:MM). Experiments performed n ≥ 3 times.

Video S8: Time-lapse video showing adhesion and spread of resOVCAR-3 RFP clusters on top of untransformed human coelomic mesothelial cells (MeT-5A). Total video duration – 44 hrs (HH:MM). Experiments performed n ≥ 3 times.
